# Asking the Data: Conversational Interrogation of LC–MS/MS Proteomics Data Sets with an LLM Agent

**DOI:** 10.64898/2026.10.22.760949

**Authors:** Carolaine S. Araújo, Milan Keirsebelik, Ana Caroline Silas Rodrigues, Alessandra L. Valverde, Rafael Maciel-de-Freitas, Marília Zaluar P. Guimarães, Magno Junqueira

## Abstract

Proteomics workflows are designed to answer predefined questions, leaving analytical problems that emerge after first results difficult to investigate without additional bioinformatics work. We introduce data set interrogation, a conversational framework in which a researcher uses a large language model (LLM) agent to translate emerging questions into executable tests, check the answers against controls, and keep the code, inputs, and decisions in a dated record. Applied retrospectively to four LC–MS/MS data sets, the framework exposed problems that changed the interpretation of three in-house studies: a search database lacking one of two symbionts, a swapped pair of tandem mass tag (TMT) channels, statistics inflated by non-host proteins, and false deamidation assignments caused by selection of the ^13^C isotope peak as monoisotopic; the same question asked of every data set revealed failed or partial cysteine alkylation in all three. A public TMT data set was reanalyzed from raw files in 24 min, a feasibility measure without a human comparison; among spectra identified by both workflows, 99.2% received the same peptide assignment as in Proteome Discoverer, and protein fold changes correlated (r = 0.93). In seeded tests, a purpose-built channel-assignment check detected all 216 between-strain channel swaps and none of 90 within-strain swaps. The framework couples natural-language interaction with executable analyses, explicit controls, and traceable decisions, enabling researchers to investigate questions not anticipated by the original pipeline. These results establish feasibility in a limited retrospective setting with one researcher; broader evaluation across users, models, and data sets is required to determine general performance.

## INTRODUCTION

The bottleneck in bottom-up proteomics has moved from acquisition to analysis. A single Orbitrap Astral injection now yields tens of thousands of MS2 spectra, and a typical study passes through format conversion, database construction, database search, rescoring, protein inference, quantification, normalization, statistics, and functional interpretation. Each step has parameters, and many choices that change the biological conclusion are never written down: which proteome version was used, which channels were assigned to which sample, which proteins entered the empirical Bayes prior, and what happened to the spectra that were not identified.

Fixed pipelines solve part of this problem. They make a defined workflow repeatable, but they answer only the questions they were built to answer. In our experience, the most consequential findings in a data set come from questions asked after the first result: why a protein that should be exclusive to one group appears in another channel, why three-quarters of the spectra are unexplained, or whether the instrument is picking the right monoisotopic peak. Answering such questions usually requires a bioinformatician, new code, and days of turnaround, so most of them are never asked.

Data already acquired therefore often stay partly unexplored once the first pipeline run has returned an answer, what is sometimes called dark data; we use the term descriptively. A misconfigured search database, a mislabeled TMT channel, or a modification call that is an instrument artifact can sit in such data unnoticed. A second motivation is traceability, the possibility of reconstructing how an analytical decision was reached. When every reported number comes from code run in the analysis session and each version of the workflow is recorded with its date and the reason for each change, a reviewer or coauthor can follow the path from the raw measurement to the reported result and tell a change of method from a change in the data. Traceability is not reproducibility, and recording a decision does not prevent an error; it makes the decision visible and open to challenge.

Large language models (LLMs) have evolved from text predictors into agents that plan, call tools, and correct themselves.^1,2^ In proteomics, an agent built a complete differential-analysis application from four natural-language prompts,^3^ and agents that retrieve published omics data and requantify them with containerized tools, under human guidance, reproduced five public studies with per-sample correlations of 0.85–0.997 to the deposited results.^4^ Hybrid workflows also use LLMs to interpret omics results mechanistically.^5^ The same literature warns about unstable reasoning, hallucination, and irreproducibility. In a randomized study of research reproduction, AI-assisted teams matched human-only teams, whereas AI-led teams with minimal oversight reproduced only 37% of results.^6^

These reports point to a design question rather than a capability question: under what working arrangement does an agent improve a proteomics analysis instead of degrading it? Here we report a use different from building an application or rerunning a published analysis. A domain expert and an LLM agent (Claude, Anthropic) worked in a conversation with a Linux workspace and analyzed data sets end to end, from RAW files to manuscripts. The expert asked questions, the agent wrote and ran the code, and every answer came back as numbers, tables, or figures that could be checked and rerun. We call this mode of work data set interrogation. Neither the use of an LLM to write analysis code nor agent-driven reanalysis is new, and we do not claim that conversation or code generation is new in itself. Our claim is narrower: in conversation, a researcher can formulate and test analytical questions that the initial pipeline did not anticipate, and in these studies such questions revealed identification, quantification, and sample-quality problems that changed what the studies could claim. The two closest systems are compared with this work after the Limitations.

Operationally, we call an analysis data set interrogation when three conditions hold: the question arises from a result rather than from a plan made before the analysis; the agent turns it into a computational test written for that question, with a control where one exists; and the question, code, inputs, answer, and the decision taken are kept in a dated record. An analyst exploring data with scripts, with or without LLM-written code, can meet the first two conditions; what the arrangement changes is that each test costs one message in a conversation, and that the record is part of the working rules rather than an afterthought. The novelty we claim lies in turning unanticipated scientific doubts into verifiable tests while preserving the history of decisions and results, not in the agent architecture or in any single module. Throughout, “agent” denotes the LLM with its tools in the workspace; “pipeline”, the fixed laboratory workflow from RAW files to statistics; and “check”, a script written to answer one question against a control.

We describe (i) the architecture of the interaction between the researcher, the agent, and the analysis tools; (ii) a benchmark of execution time, cost, human intervention, reliability, and reproducibility; (iii) three case studies, on building and auditing search databases, on quality control of peptide-spectrum matches (PSMs), and on proteomics of non-model organisms; (iv) validation against established software, reference methods, and independent controls; and (v) the limitations, including errors made by the agent, the effect of incorrect metadata, and the dependence on the researcher’s expertise. The case studies come from three in-house studies whose biological results are reported in companion studies;^7−9^ here we report only what is needed to evaluate the analysis. A public data set that any reader can reanalyze serves as the reproducible case. The three in-house data sets were chosen to differ: TMTpro-labeled neurospheres derived from human induced pluripotent stem cells modeling a monogenic disease, TMTpro-labeled insect ovaries and heads under symbiont infection and two thermal regimes, and a label-free proteome of a non-model marine holobiont compared across preparation workflows and specimens. This diversity probes whether the approach is useful beyond one organism, design, or class of problem; it is not by itself evidence of broad applicability.

## EXPERIMENTAL SECTION

### Working Environment

All computation ran in a private Linux workspace attached to a Claude conversation (claude.ai, Project mode), with a shell, Python, file tools, about 8 GiB of RAM (7.8 GiB measured on 14 October 2026), two CPU cores, and no GPU. Files on the corresponding author’s computer were reached through the desktop application, which copies selected files into the workspace and writes results back. The corresponding author interacted with the agent only in natural language, in Portuguese. The analyses ran in Claude models available in claude.ai in October 2026; the sessions of 13 and 14 October, which ran the revision analyses and the public-data reanalysis, used Claude Opus 5.5; earlier sessions used the Claude models then available in claude.ai. Four conditions of execution can change the results and are reported for that reason: the model version; the context stored in the account, whose memory index reached every session (Limitations); web access, which was allowed for searches while UniProt and PRIDE could not be reached from the shell; and the workspace resources.

The analyses ran in a consumer claude.ai account. Under the provider’s consumer terms, conversations are used for model training only when the user enables the model training setting, and data are kept for 30 days when it is off.^10^ Raw files from the in-house studies were processed in the workspace with the agreement of the data owners, who are authors of this work; the iPSC-derived data contain no donor-identifying information.

### Data Sets

Three in-house studies and one public data set provided the data (Table 1). In the mosquito study, *Aedes aegypti* lines carrying either *w*Mel or *w*AlbB, two strains of the maternally inherited endosymbiont *Wolbachia*, and backcrossed to a local uninfected genetic background (URCA),^11^ were reared under two thermal regimes (18–26 °C or 26–34 °C); ovaries and heads from three biological replicates were labeled with TMTpro 18plex (Thermo Fisher Scientific, Waltham, MA) using the miniaturized Opt-TMT protocol^12^ and analyzed by single-shot injections (200 ng) on an Orbitrap Astral. The same spectra also measured the insect-specific Phasi Charoen-like virus (PCLV), which persists in Ae. aegypti cell lines (Case Study 3). In the Dravet study, neurospheres derived from induced pluripotent stem cells of a control line (GM23279A) and a Dravet syndrome line (DRV3, carrying *SCN1A* and *SCN9A* variants) were kept at 37 °C or exposed to 42 °C for 30 min followed by 3 h of recovery, prepared as described,^12,13^ labeled with TMTpro 16plex (12 channels used), and acquired on an Orbitrap Exploris 480 and an Orbitrap Astral. In the sponge study, *Viles ophiraphidites* (formerly *Topsentia ophiraphidites*) specimens were analyzed label-free on a Q Exactive, first comparing four preparation workflows (42 runs) and then four specimens prepared by S-Trap in technical triplicate (12 runs).

**Table 1.** Data Sets.

| Data set | Design | Instrument | Injections | Main question |
| --- | --- | --- | --- | --- |
| *A. aegypti* × *Wolbachia*<sup>7</sup> | TMTpro 18plex; 3 strains × 2 thermal regimes × 3 biological replicates; ovary and head | Orbitrap Astral | 6 (3 per tissue); one further 300-ng head injection excluded | Does infection or heat remodel the host proteome? |
| Dravet neurospheres<sup>8</sup> | TMTpro 16plex (12 channels); GM23279A vs DRV3; 37 vs 42 °C | Orbitrap Exploris 480; Orbitrap Astral | 4 | Genotype and hyperthermia effects; reliability of identifications |
| *V. ophiraphidites*<sup>9</sup> | Label-free; 4 preparation workflows (phase 1); 4 specimens by S-Trap (phase 2) | Q Exactive | 54 | First proteome of a non-model sponge |
| NSC vs neurospheres, PXD063185 (public)<sup>14</sup> | TMT 10plex; 5 NSC vs 5 NPH; one plex injected twice | Q Exactive Plus | 2 | Reproducible case: are the deposited results recovered? |

The public data set, PXD063185, compares five two-dimensional neural stem cell cultures (NSC) with five three-dimensional neurospheres (NPH) from one hiPSC line, labeled with TMT 10plex in a single plex and injected twice as single-shot runs (technical replicates) on a Q Exactive Plus with 68 min of acquisition; the original analysis used Proteome Discoverer 3.1.1.93 with Sequest HT against the 2023 human Swiss-Prot proteome, with phosphorylation and N-terminal acetylation as additional variable modifications.^14^ The channel map (NSC_1–NSC_5 in 127N, 128N, 129N, 130N, and 131; NPH_1–NPH_5 in 126, 127C, 128C, 129C, and 130C) was taken from the original article.^14^

The three in-house data sets were generated by the authors and their colleagues for studies that are not yet published. To avoid preempting those studies, we report only results that bear on the analysis: database coverage, channel assignment, behavior of the statistics, fate of the spectra, and quality of the identifications. Biological findings, including absolute numbers of altered proteins, are reported in the companion studies.^7−9^ The use of patient-derived induced pluripotent stem cell lines was approved by the Institutional Research Ethics Committee of IDOR and by the Brazilian National Research Ethics Committee (protocol 67185123.2.0000.5249), and written informed consent was obtained from the patient’s legal guardians before sample collection, as described for the same cohort.^15^ No unexpected or unusually high safety hazards were encountered.

### Software and Databases

RAW files were converted to mzML or MGF with ThermoRawFileParser (2.0.0-dev)^16^ using vendor centroids and instrument-assigned charges. Database searches used Sage 0.14.6 and 0.14.7^17^ with reversed-sequence decoys and 1% FDR, with isotope errors 0 and 1 unless stated otherwise (−1 to +1 in the metaproteomic search; −1 to +3 in FragPipe); fragment annotation for the envelope audit used Sage 0.15 (--annotate-matches). FragPipe 24 (MSFragger 4.4.1,^18^ Percolator, Philosopher/ProteinProphet, IonQuant, TMT-Integrator) served as the second engine. Open searches of the mosquito data used Sage with precursor_tol set to −400 to +150 Da, which Sage defines as theoretical minus experimental mass; the search therefore accepted mass shifts (experimental minus theoretical) from −150 to +400 Da. The open search for cysteine forms in the Dravet data accepted mass shifts from −150 to +350 Da. Unexplained spectra were sequenced de novo with an in-house spectrum-graph algorithm (denovo.py; b/y prefix-mass nodes scored by ion evidence, best path by dynamic programming, precursor masses M and M − 1.00335 tested), and sequences were matched by MS-BLAST-style alignment with a PAM30MS matrix against *Drosophila melanogaster* (UP000000803) for the mosquito data and against *Halichondria panicea* for the sponge data.

Databases were built through the UniProt REST API (release 2026_05) with documented proteome identifiers, genome assemblies, and BUSCO completeness. The mosquito database combined *A. aegypti* LVP_AGWG (UP000008820; 20,643 entries), *w*Mel (UP000008215; 1,166), *w*AlbB (nonredundant union of UP000290937 and UP000004822; 1,295), and 379 universal contaminants,^19^ for 23,483 entries without decoys. The sponge databases combined sponge and symbiont sequences with the proteome of H. panicea, the closest relative with a sequenced genome: DB-2 (497,112 targets; MD5 51d45b13b2fbfb4c726704b0d2164775) for phase 1 and the agent sessions, and DB-3 for phase 2 (DB-2 without Vibrio entries and with an updated contaminant list; 466,314 targets and 429 contaminants; MD5 c392944394ee74f2448fa0bacdff7943). A metaproteomic database of 13 bacteria, 2 yeasts, and *Aedes*-associated viruses (77,015 proteins) was built for the unexplained spectra. The human database for PXD063185 was the reviewed human proteome (UP000005640, Swiss-Prot canonical, 20,416 entries, downloaded on 9 October 2026) with 49 contaminants.

### TMT Quantification and Statistics

For the mosquito study, reporter ions were extracted within ±3 mDa from spectra with precursor isolation purity ≥0.5 and corrected for lot-specific isotopic impurities by nonnegative least squares (NNLS). Only PSMs unique to host or to *Wolbachia* were used. Reporter intensities were summed per protein over PSMs, log2-transformed, median-normalized per channel, centered within each injection, and averaged over the three injections of each sample; host channels were then renormalized to the median of host proteins only. For the mosquito study, a linear model with strain × regime and biological replicate as a block was fitted for each protein, followed by empirical Bayes moderation^20,21^ and Benjamini–Hochberg correction, both computed on host proteins only; 13 host proteins whose signal was restricted to infected channels were excluded from the ovary model (Table S2). The first analysis version (v1) had instead included *Wolbachia* and contaminant proteins in normalization, prior, and correction. Robustness was checked with quantile normalization, and minimum detectable effects at 80% power were computed from the residual variance. PCLV reporter intensities were read from the same MS2 scans, divided by each channel’s summed host reporter signal, and expressed per PSM as channel share; groups were compared by two-sided t-tests on log2 values, as in the companion study.^7^

### Reanalysis of PXD063185

The two RAW files were located on the corresponding author’s computer by the agent, copied to the workspace, and processed with the laboratory’s pipeline as written by the agent in earlier sessions (Supporting Information): ThermoRawFileParser 2.0.0-dev; Sage (v0.14.7 release; the binary reports version 0.14.6) against the human database with reversed decoys; trypsin, up to 2 missed cleavages, 7–30 residues; carbamidomethyl-Cys and TMT (+229.1629 Da on Lys and peptide N-terminus) fixed; Met oxidation and protein N-terminal acetylation variable (up to 3); precursor ±10 ppm, fragment ±0.02 Da, isotope errors 0 and 1; 1% PSM and protein FDR with parsimony. Reporters were extracted from vendor centroids (±3 mDa) in spectra with isolation purity ≥0.5 and were not corrected for impurities, because the lot certificate was not available. Protein abundance was computed as the sum of the reporters of razor spectra, log_2_-transformed, median-normalized per channel, centered for each protein within each injection, and averaged over the two injections. As in the original analysis, NPH and NSC were compared by a two-sample t-test at p < 0.01, without correction for multiple testing; this test was used only to match the original report, whereas the in-house studies used moderated statistics with Benjamini–Hochberg correction, so counts of significant proteins are not comparable across data sets. The deposited mzTab file (Proteome Discoverer 3, Sequest HT) was parsed for PSMs by run and scan and for protein abundances by assay; its abundances were log_2_-transformed, median-subtracted per assay, averaged over injections, and tested in the same way. Spectra were matched by run and scan, and peptides were compared without modifications and with I = L. The human database used here (2026) was newer than the original one (2023). Two further searches tested the comparison. In the first, phosphorylation of S, T, and Y was added as a variable modification (up to two per peptide), with the database split into six parts to fit in memory and FDR computed globally. In the second, an entrapment search, every human protein was paired with a shuffled copy (residues shuffled within each tryptic peptide, C-terminal K or R kept), so that matches to the shuffled copies estimate the false discovery proportion by the combined method with an entrapment-to-target ratio of 1.

### Fragment Tolerance Test

The Orbitrap Exploris 480 run of the Dravet data set was searched with Sage (v0.14.7 release; the binary reports version 0.14.6) at fragment tolerances of ±20, ±10, and ±5 ppm with otherwise identical parameters (TMTpro fixed on Lys and peptide N-terminus, human database). PSMs were filtered at 1% FDR by target-decoy competition on the hyperscore, because Sage’s linear rescoring failed and fell back to a heuristic score in two of the three searches, which made its q-values incomparable across settings.

### Reproducibility of the Agent

Three questions from this work, two on PXD063185 and one on the mosquito study, were re-asked, each in two new agent sessions started without the conversation history and instructed not to open the original scripts or results: the peptide agreement between Sage and the deposited Proteome Discoverer results for PXD063185; which sample fitted its own group worst in PXD063185; and the number of host proteins altered by *w*Mel in the mosquito ovary when the prior and correction are computed on host proteins only or on all proteins. Each session received the question and the input files, wrote its own code, and reported numbers that were then compared with this manuscript.

### Simulated Non-Specialist Requests

To test how the wording of a request changes the analysis, PXD063185 was given to new agent sessions under four scenarios, three sessions each (Table S11). Each session started in an empty folder with the two RAW files, offline human Swiss-Prot and cRAP databases, ThermoRawFileParser, and Sage, with web search allowed but UniProt and PRIDE unreachable from the workspace, and received a request written as a cell biologist without proteomics training would write it: (A) a vague request naming only the two culture conditions; (B) the same request with the facility’s e-mail, which gives the TMT10 channel map and states that the files are two injections of one pool; (C) a request with a wrong premise, that each file holds one condition; or (D) the expert request with all parameters. A second agent played the non-specialist from a fixed character sheet: it could paste the facility’s e-mail only when asked about labeling, channels, or which sample was which, and answered that it did not know anything else; questions in scenario D were answered from the request. A session ended when results were delivered. Every tool call was audited for file paths outside the session folder, web searches, and use of the corresponding author’s stored memory. Results were compared with the validated pipeline result (moderated test with Benjamini–Hochberg correction: 1,393 proteins tested, 607 at q < 0.05; Table S8a) and with the deposited Proteome Discoverer results. Twelve of the first 16 sessions (which included four exploratory sessions without web access) read the stored memory, which names PXD063185 and its design, despite an instruction not to; all 12 were excluded, and the eight that belonged to the main design were repeated with all memory and connector tools forbidden. Every session nonetheless received an automatic index of that memory, which could not be switched off in the workspace. The sponge sessions described below were audited in the same way, for 30 sessions in all; one sponge session was interrupted by the user before it delivered results and is therefore audited but not listed in Table S11. The same design was applied to three technical replicate injections of the sponge, with sessions of an agent type that had no memory tools: non-specialist sessions received the RAW files and a request asking which proteins and bacteria were present, and were given the sponge database only if they asked for one; the expert session received the database and the parameters of the laboratory pipeline. Alkylation was checked by searching one injection from each phase against the *Halichondria panicea* proteome with carbamidomethylation of cysteine as a variable modification, and then in all 54 sponge runs with Sage against the 6,001 proteins identified in 12 representative runs and their reversed decoys, with no fixed modification and with cysteine carbamidomethylation, disulfide bonding (−1.0078 Da), dioxidation, trioxidation and seven other cysteine adducts (propionamide, carboxymethyl, N-ethylmaleimide, pyridylethyl, methylthio and two others) as variable modifications (Table S12).

### Benchmark Measures

Execution time for PXD063185 was taken from the timestamps of the corresponding author’s message, the copied files, and the pipeline logs. Cost was taken from the subscription invoices. Human interventions were counted as messages from the corresponding author that started or redirected an analysis and as decisions about samples taken with the data owners, from the conversation and the dated reports (Table S1). Reproducibility was tested by rerunning the complete pipeline on one PXD063185 injection from the RAW file in a new directory and comparing MD5 checksums of the PSM, protein-group, reporter, de novo, and similarity tables, and by regenerating every figure and number in this manuscript from scripts and source data. Time invested by the corresponding author was estimated from the modification times of the files in his download folder, and in the folders the agent wrote to on his computer and in his documents folder, from the first file of this work (sponge database and first filtered sponge MGF file, 9 October 2026, 00:21) to the final version of the sponge companion manuscript (20 October 2026, 08:00). Files were assigned to stages by name; versions of this manuscript, the coauthor versions with their marked reviews and suggestions, the responses to pre-submission critiques, and the companion manuscripts were assigned to their own stages; files of unrelated activities, teaching, administration, conference slides, and a fourth data set not reported here were excluded. Files saved within 60 min of each other were merged into one working session, each counted from 15 min before its first file to its last file; gaps of 45 and 90 min gave the range (Table S10). The estimate counts only periods in which files were saved and is a lower bound.

### Seeded-Error Test and Null Test

To test the checks against a standard workflow, we introduced known defects into the mosquito data (Tables S5–S7). (1) Every possible swap of two channels (153 per tissue: 108 between channels of different strains, 36 for each pair of strains, and 45 within a strain; 306 in both tissues) was applied to the protein-level matrices and to the strain-specific peptide signal. Standard sample QC assigned each channel to the group whose leave-one-out centroid correlated best with it over the 500 most variable host proteins; ChannelGuard assigned channels by strain-specific peptide signal with group quotas taken from the design. Peptides unique to *w*Mel or to *w*AlbB were used, rather than a pooled *W. pipientis* database, because only strain-specific peptides can tell the two infected groups apart. A swap counted as detected when a swapped channel was flagged and had not been flagged on the true map. (2) For the statistics, host channels were permuted within each thermal regime 200 times, which removes any strain effect in the host while symbiont and contaminant rows keep their true channels; every host protein called at *q* < 0.05 is then a false positive. Four pipelines were compared: host-only normalization, prior, and correction (v2); all-protein prior and correction; all-protein normalization; and all three on all proteins (v1). (3) For databases, DBScope audited the correct database, the inherited file, and versions with 10%, 50%, or 100% of *w*AlbB, the contaminants, or the host proteome removed, flagging any reference proteome with coverage below 90%. The ChannelGuard assignment rule was revised once after its first version flagged one correct ovary channel; the head data were then evaluated without further change.

### Verification of Isotopic Envelopes

For every PSM at 1% FDR, each annotated fragment and each precursor was compared with the vendor centroids in the corresponding scan. Theoretical envelopes were computed from the exact elemental composition, including the heavy atoms of TMTpro (7 ^13^C and 2 ^15^N per label; enrichment 99.42%), and conditioned on the precursor isolation window (0.5 *m*/*z* nominal; effective half-window 0.32–0.35 *m*/*z*). Fragments were tested for an M+1 peak at Δ/*z* with the expected ratio, for a peak at −Δ/*z* that would make the assigned peak an M+1 of another ion, and for the spacing of another charge; offsets with no isotopic meaning served as negative controls. Precursors were tested for the isotope selected by the instrument, cosine similarity to the theoretical envelope, a shifted monoisotopic peak, and a missing monoisotopic peak. The fraction of random fragment matches was estimated with a mixture of the error distribution of coherent fragments and a uniform distribution over the ±20 ppm tolerance.

### Combination of Identifications

Sage and FragPipe results were combined at a global 1% FDR across all runs, and protein groups were inferred by parsimony on the union of peptides (I = L). A de novo sequence was considered concordant with a PSM when the two shared at least four consecutive residues; the rate of this criterion on random spectra was 0.2%.

### Data and Code Availability

Raw files and search results of the sponge study are deposited to the ProteomeXchange Consortium via PRIDE (PXD079594, with phase 2 being added), and those of the mosquito and Dravet studies are being deposited; the three data sets are described in the companion studies7−9 and will be made public with their publication. The public case uses PXD063185. The Interroga package (tested modules, the laboratory pipeline, scripts as run, and every script that regenerates the figures and analyses from source data) is provided as Supporting Information and will be released in a public repository. Table S4 states for each component whether it was run during the analyses and released as run, refactored from those scripts with unit tests added, written after the analyses (DogWatch), or used in the analyses and available from the corresponding author on request (FragSlice, Penumbra, and the Dravet combination script), and Table S14 maps each central result to the script and input files that produce it. The interactive page for the sponge data set is available from the corresponding author on request.

## RESULTS AND DISCUSSION

### Architecture: How the Researcher, the Agent, and the Tools Interact

The work ran as a loop with three participants (Figure 1a). The researcher asks a question in plain language. The agent translates it into code, runs the code in the workspace against the data, tests the result against an internal control when one exists, and reports numbers, tables, or figures. The researcher accepts the answer, asks a follow-up question or, when samples are involved, decides together with the data owners; the accepted result is then frozen in a dated report. The analysis tools sit in the workspace (Figure 1b): format conversion, search engines, de novo sequencing, homology search, quantification, and statistics form the standard pipeline, and new modules are added when a question requires one. The record of the work is not the conversation but what the loop leaves behind: scripts, versioned reports, and source data for every figure. Durable context, such as database versions, sample maps, decisions, and status, was written to Markdown reports stored in the claude.ai Project, so that later sessions started from the documented state rather than from the chat history.

**Figure 1.**
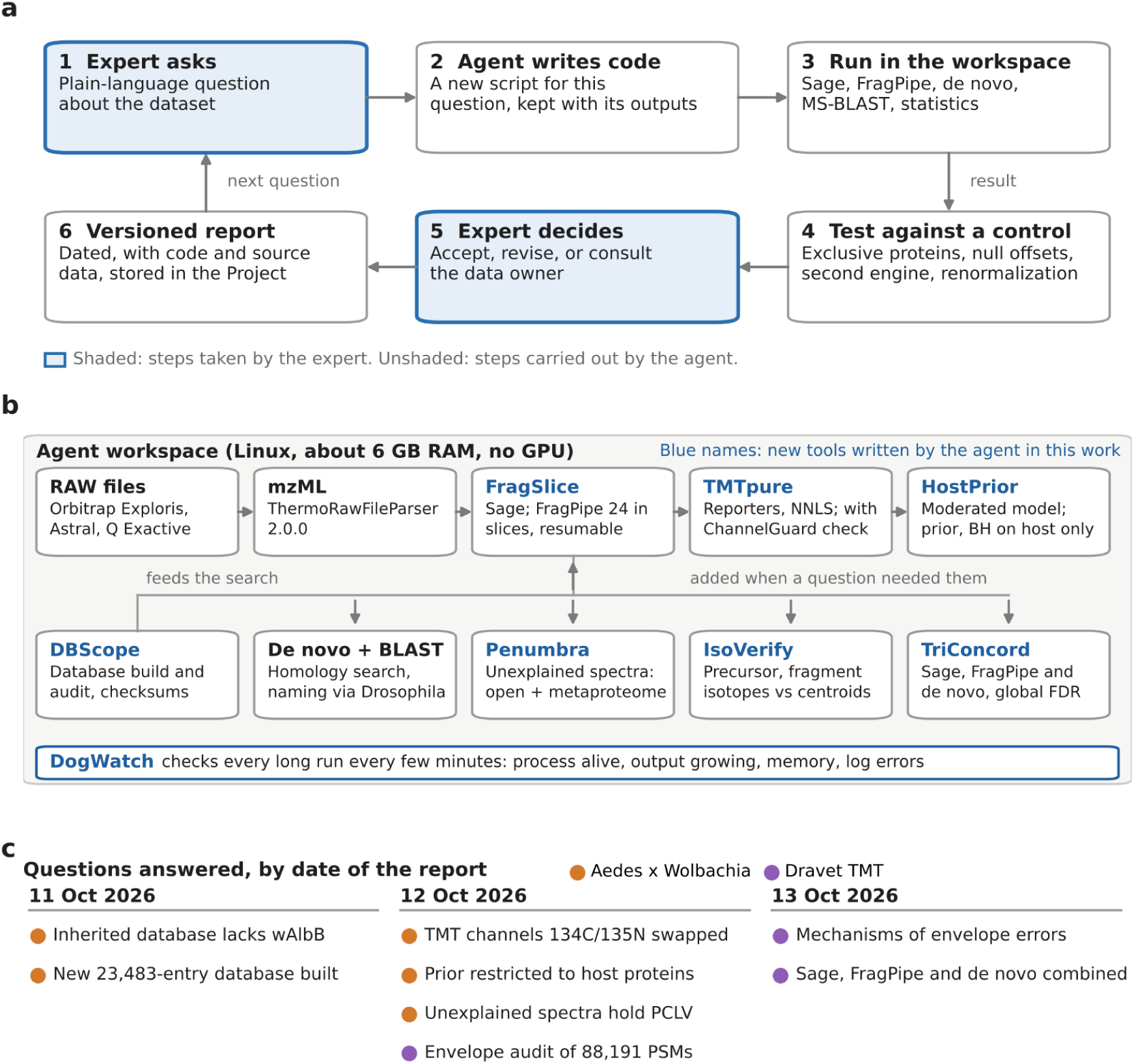
Architecture of data set interrogation with an LLM agent. (a) The loop used in all studies. The researcher asks a question in plain language; the agent writes and runs new code, tests the result against an internal control, and reports it; the researcher accepts, revises, or consults the data owner; the result is frozen in a dated report. Shaded boxes are the researcher’s steps. (b) Modules in the agent’s workspace. The standard pipeline (top row) runs from RAW files to statistics; the modules in the bottom row were added because a question required them. Names in blue are checks written by the agent in this work. The strip at the bottom is DogWatch, which supervises long runs; during the analyses it was written anew whenever the researcher asked for it. (c) Questions answered in conversation between 11 and 13 October 2026, by date of the dated report and by study.

Five rules governed the loop. (1) Every number in a report came from code that the agent ran in that session, never from recall. (2) Each analysis version was frozen as a dated report (v1, v2, v3), and changes between versions were stated with their reason. (3) Claims were tested against an internal control whenever one existed: proteins exclusive to *Wolbachia*, offsets with no isotopic meaning, shuffled sequences, a second search engine, or an alternative normalization. (4) Decisions about samples, such as channel reassignment or exclusion of an injection, were proposed by the agent with evidence and taken by the researcher together with the data owner. (5) Code, intermediate tables, and source data for every figure were kept and exported with the reports.

Eight checks were written by the agent during this work, each in response to a specific question, between 11 and 13 October 2026 (Table 2). Most implement established methods, such as NNLS impurity correction or empirical Bayes moderation, and we do not claim methodological novelty for them. What we report is that each was prompted by a concrete problem in real data, and how each performed. Four core modules (DBScope, ChannelGuard, TMTpure, HostPrior) and the run supervisor DogWatch, described below, are released as a tested Python package under the name Interroga; IsoVerify is released as the scripts that were run; FragSlice, Penumbra, and the Dravet part of TriConcord were used in the analyses and are available from the corresponding author on request. EagleEye, an MGF filter developed earlier in our laboratory, predates this work and is not part of the set. Table 3 lists the questions that had the largest effect; none of them belonged to the pipeline as first designed.

**Table 2.** Checks Written by the Agent during This Work.

| Tool | Function | Question that created it | Validation and status |
| --- | --- | --- | --- |
| DBScope | Builds search databases from UniProt proteomes with assembly, BUSCO, and checksum records; removes redundancy; adds contaminants; audits an existing database by sequence identity to reference proteomes | Does the inherited database cover both symbionts? | Seeded database defects; unit tests; released |
| ChannelGuard | Checks each TMT channel against the signal of peptides exclusive to one group and flags swapped channels | Are the ovary channels assigned correctly? | 306 seeded channel swaps; unit tests; released |
| TMTpure | Extracts TMTpro reporters ($\pm 3$ mDa), filters by isolation purity ($\geq 0.5$), and corrects lot-specific impurities by NNLS | How are reporters quantified without vendor software? | Recovers synthetic intensities (unit test); comparison with vendor correction planned; released |
| HostPrior | Moderated linear model with block effects in which the variance prior and Benjamini–Hochberg correction use host proteins only; quantile-normalization robustness and minimum detectable effect | Why do so many host proteins change? | Independent reimplementations; permutation null test; unit tests; released |
| FragSlice | Runs FragPipe in about 6 GB of RAM by slicing the database and mzML files; resumes from completed slices and relaunches after interruptions | Can a second engine run in this environment? | Used for the second-engine search; available from the corresponding author on request |
| Penumbra | Accounts for the fate of every MS2 spectrum; runs open search and a metaproteomic search with class-specific FDR; removes shared peptides; quantifies microbial peptides by TMT | What are the 72–76% unexplained spectra? | Used for the unexplained spectra; available from the corresponding author on request |
| IsoVerify | Verifies precursor and fragment isotopic envelopes against vendor centroids, including TMTpro heavy atoms and the isolation window, with null-offset controls | Is the instrument choosing the right monoisotopic peak? | Null-offset controls; scripts as run released |
| TriConcord | Combines Sage, FragPipe, and de novo results at a global FDR; infers protein groups by parsimony; assigns evidence levels; exports a verified spectrum set | Do two engines agree, and which spectra can be trusted? | Parsimony and FDR merge released; combination script available from the corresponding author on request |

**Table 3.**
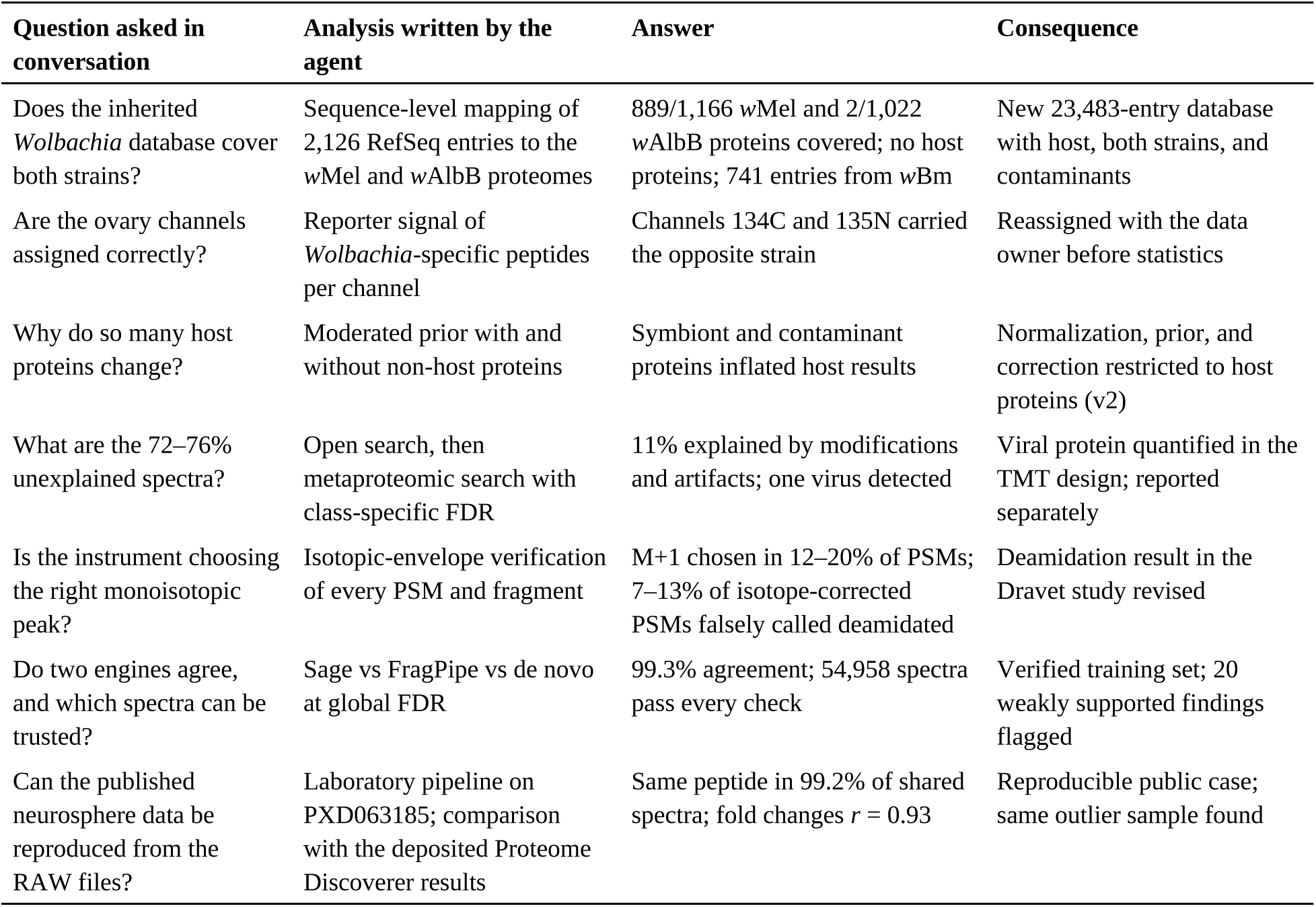
Questions Posed to the Data Sets and Their Consequences.

| Question asked in conversation | Analysis written by the agent | Answer | Consequence |
| --- | --- | --- | --- |
| Does the inherited *Wolbachia* database cover both strains? | Sequence-level mapping of 2,126 RefSeq entries to the wMel and wAlbB proteomes | 889/1,166 wMel and 2/1,022 wAlbB proteins covered; no host proteins; 741 entries from wBm | New 23,483-entry database with host, both strains, and contaminants |
| Are the ovary channels assigned correctly? | Reporter signal of *Wolbachia*-specific peptides per channel | Channels 134C and 135N carried the opposite strain | Reassigned with the data owner before statistics |
| Why do so many host proteins change? | Moderated prior with and without non-host proteins | Symbiont and contaminant proteins inflated host results | Normalization, prior, and correction restricted to host proteins (v2) |
| What are the 72–76% unexplained spectra? | Open search, then metaproteomic search with class-specific FDR | 11% explained by modifications and artifacts; one virus detected | Viral protein quantified in the TMT design; reported separately |
| Is the instrument choosing the right monoisotopic peak? | Isotopic-envelope verification of every PSM and fragment | M+1 chosen in 12–20% of PSMs; 7–13% of isotope-corrected PSMs falsely called deamidated | Deamidation result in the Dravet study revised |
| Do two engines agree, and which spectra can be trusted? | Sage vs FragPipe vs de novo at global FDR | 99.3% agreement; 54,958 spectra pass every check | Verified training set; 20 weakly supported findings flagged |
| Can the published neurosphere data be reproduced from the RAW files? | Laboratory pipeline on PXD063185; comparison with the deposited Proteome Discoverer results | Same peptide in 99.2% of shared spectra; fold changes $r = 0.93$ | Reproducible public case; same outlier sample found |

#### Supervising runs: DogWatch

The tool that saved the most time was not a check on the data. Searches of Astral files, of sliced databases and with FragPipe ran for tens of minutes to hours in a workspace with about 8 GiB of RAM, where jobs were killed by memory limits and sessions were interrupted. The agent did not watch these runs on its own. Throughout the case studies, the corresponding author had to ask it again and again to “dog watch the runs”; each time, the agent wrote a new check that every few minutes tested whether the process was alive, whether its output was growing, how much memory it used and whether the log reported an error, and relaunched the job when it could be resumed (an early version, vigia.sh, relaunched FragPipe slices after an interruption). Because each check was written on request, it differed between sessions and was not kept. The sponge sessions of the non-specialist test show the same behavior with no person present: in sessions of 43–325 min they checked the state of their jobs 7–28 times each and waited a median of 100–420 s between checks, never more than 10 min, the longest a single tool call can last; waiting took 23–129 min per session. A check runs only while the agent is working on a turn. When the turn ends or the session is interrupted, the job continues unwatched until someone asks; when the workspace restarted during one of these sessions, the agent established which outputs had survived only after it was resumed. We therefore consolidated the checks into DogWatch (interroga/dogwatch.py, with tests), which supervises a process or a command, records each check as running, stalled, finished or failed in a status file that a later session or a person can read, and can relaunch a resumable job when the run stalls or fails (Figure 1b). This version was written after the analyses reported here; during them, supervision of the runs depended on the researcher asking for it.

### Benchmark: Time, Cost, Human Intervention, Reliability, and Reproducibility

Table 4 summarizes the benchmark. The public data set gave the cleanest timing. From the corresponding author’s one-line request to find the RAW files on his computer to a validated comparison with the deposited results took 24 min of wall-clock time. This is a feasibility measure for one small data set: no analyst performed the same task for comparison, so it does not show that the approach is faster than conventional analysis. Copying the two RAW files (425 MB) took about 10 min, installing the converter and search engine about 3 min, and the pipeline 10.2 min of wall-clock time, of which 6.4 min were computation: 3.0 and 3.2 min per injection from RAW file to quantified proteins, including de novo sequencing and homology search, and 14 s for the comparison between conditions; the rest was spent fixing a step of the pipeline that stopped once (see Limitations). The comparison with the deposited results took under 1 min. The in-house studies were not timed as systematically: their 10 main questions (Table S1) were answered over three days, each within the session in which it was asked.

**Table 4.** Benchmark of Data Set Interrogation.

| Measure | Result | Source |
| --- | --- | --- |
| Execution time | PXD063185 (2 RAW files, 425 MB): 24 min from request to validated comparison; 6.4 min of pipeline computation (3.0 and 3.2 min per injection; 14 s for the comparison) | Message, file, and log timestamps |
| Cost | Consumer subscription: Claude Pro (R\$110 per month) until 11 October 2026, then Max 5x (R\$550 per month); no per-token, cloud, or software license charges; the subscription also served other work | Invoices |
| Researcher time | About 50 h of active work (43–61 h) in 50 sessions over 11 days and 8 h, from the first sponge file to the final companion manuscript, including revision with coauthors; 28 h (26–32 h) to the first complete draft; stages in Table S10 | File timestamps on the researcher's computer |
| Computing resources | 7.8 GiB of RAM (measured on 14 October 2026), 2 CPU cores, no GPU; FragPipe and the phosphopeptide search required database slicing | Workspace |
| Human intervention | PXD063185: 1 instruction, no correction. In-house studies: 10 questions that started new analyses; 2 decisions about samples (one channel reassignment, one injection exclusion) taken with the data owners; 5 of 18 recorded problems found because a person asked or corrected | Conversation; Tables S1 and S3 |
| Reliability | Same peptide as Proteome Discoverer in 99.2% and as FragPipe in 99.3% of shared spectra; all 216 seeded between-strain channel swaps detected (95% CI 98.3–100%) and none of 90 within-strain swaps; at least one false host call in 3% of null runs (6 of 200; 1.1–6.4%) | Validation sections |
| Reproducibility | Complete rerun of one injection from RAW gave byte-identical PSM, protein-group, reporter, de novo, and similarity tables; three questions re-asked in six independent agent sessions of one model version (Claude Opus 5.5) returned the same answers (preliminary: three questions, one model); every figure regenerated from scripts | MD5 checksums; agent sessions (Table S8) |

#### Time invested by the researcher

The work began on 9 October 2026 at 00:21, when the sponge database and the first filtered sponge MGF file were saved (the sponge RAW files had been downloaded on 6 July for other purposes); the last file counted here, the final version of the sponge companion manuscript, was saved on 20 October at 08:00, 11 days and 8 h later. File timestamps place about 50 h of active work (43–61 h depending on the gap rule) in 50 sessions on 12 days (Table S10). By stage, with overlaps because stages ran in parallel: development and improvement of the pipeline and checks, including the Interroga package, 8.7 h (7.3–12.4); the test on the public data set and its comparison with the published results, 2.1 h (2.1–2.9); analysis of and questions to the three in-house data sets, 19.6 h (18.3–22.4; sponge 11.8, mosquito 4.9, Dravet 3.5); figures, 8.4 h (7.7–11.7); writing and revising this manuscript and its Supporting Information through 27 dated versions, 14.2 h (11.9–21.6); and the exchange with the coauthors, the students who generated the data sets and the biologists of each study, over versions of this manuscript and of the three companion manuscripts, their marked reviews and suggestions, and the responses to pre-submission critiques, 12.9 h (10.2–15.2). The first five days, up to the first complete draft of this manuscript, took 28 h (26–32 h); the following six days of revision with the coauthors took 19 h (15–25 h). These are lower bounds: reading, thinking, and conversation with the agent between saved files are not counted, nor are the meetings, laboratory discussions, and messages with the students and biologists, which raised many of the questions in Table S1. The figure describes the effort of one experienced user across a broad scope; it is not a comparison with conventional analysis. It does show where the expert’s time went: the agent shortened computation, but about a quarter of the counted time went to discussing results and versions with the people who produced the samples and the biology, and that time is part of the method.

#### Controls and conversation at every step

None of the problems in the case studies was settled by the agent alone or by the pipeline alone. Each answer became usable when it was tested against a control and then discussed with the people who knew another part of the work: the students who prepared the samples and acquired the data, who could confirm or correct a sample map or a preparation step, and the biologists of each study, who judged which results mattered and asked the next questions. Every check reported here carries its own control (offsets with no isotopic meaning, seeded defects, permutations, entrapment sequences, decoys, an alkylated reference data set), and every decision about samples was taken with the people who generated them (Table S1). The agent made each round of questions cheap; the controls made each answer checkable; the exchange between sample generation, data acquisition, analysis, and biology decided what was changed. We regard this loop, rather than the agent, as the working unit of data set interrogation, and the time it took, including the rounds of manuscript versions with coauthors, as part of the method’s cost.

The only cost was a consumer subscription that also served other work, so the cost attributable to this study is a contextual estimate rather than a measured total; nothing else was paid for software or computing. Human intervention was concentrated where it mattered: the researcher asked the questions and took decisions about samples, and the agent did not change a sample map, exclude an injection, or alter metadata on its own. Reliability and reproducibility are examined in the validation sections below. The benchmark has two clear limits. Timing comes from one small public data set; the in-house analyses involved much larger files (2–3.6 GB per Astral injection) and runs interrupted by memory limits. And the counts of human intervention describe one experienced user; they say nothing about how often a less experienced user would ask the questions that mattered.

### Case Study 1: Building and Auditing Search Databases

The first question in the mosquito study concerned the database, not the spectra. The *Wolbachia* database used in a previous analysis of the data had 2,126 RefSeq WP_ entries, including one duplicate, with no *Aedes* proteins and no contaminants. It was a generic RefSeq download: 1,385 entries were labeled MULTISPECIES (WP_ accessions group identical proteins from several genomes), and 741 occurred only in the genome of the *Wolbachia* of the filarial nematode *Brugia malayi* (*w*Bm, supergroup D). Mapping each sequence to the reference proteomes with DBScope showed that the file contained identical sequences for 889 of 1,166 *w*Mel proteins but for only 2 of the 1,022 proteins of the *w*AlbB reference proteome, both shared with *w*Mel. *w*Mel and *w*AlbB belong to the same species, *W. pipientis*, but to different supergroups (A and B); the closest matches of *w*AlbB proteins in the file had a median identity of 85%, too low for most tryptic peptides to be identical, so *w*AlbB-specific peptides could not be identified; 1,236 sequences belonged to neither strain (Figure 2). The agent then built a new database from documented UniProt proteomes, with proteome identifiers, genome assemblies, BUSCO completeness, and checksums recorded. The audit took one question and one script; without it, one of the two symbionts would have been nearly invisible to the search. With the new database, the samples yielded protein groups for 27% of the A. aegypti proteome (5,616), 31% of wMel (362), and 17% of wAlbB (222) (Figure 2a). In the seeded-error test, DBScope flagged every database variant with 50% or more of a proteome missing, including the inherited file, but not a 10% loss of *w*AlbB at the 90% threshold (see Validation).

**Figure 2.**
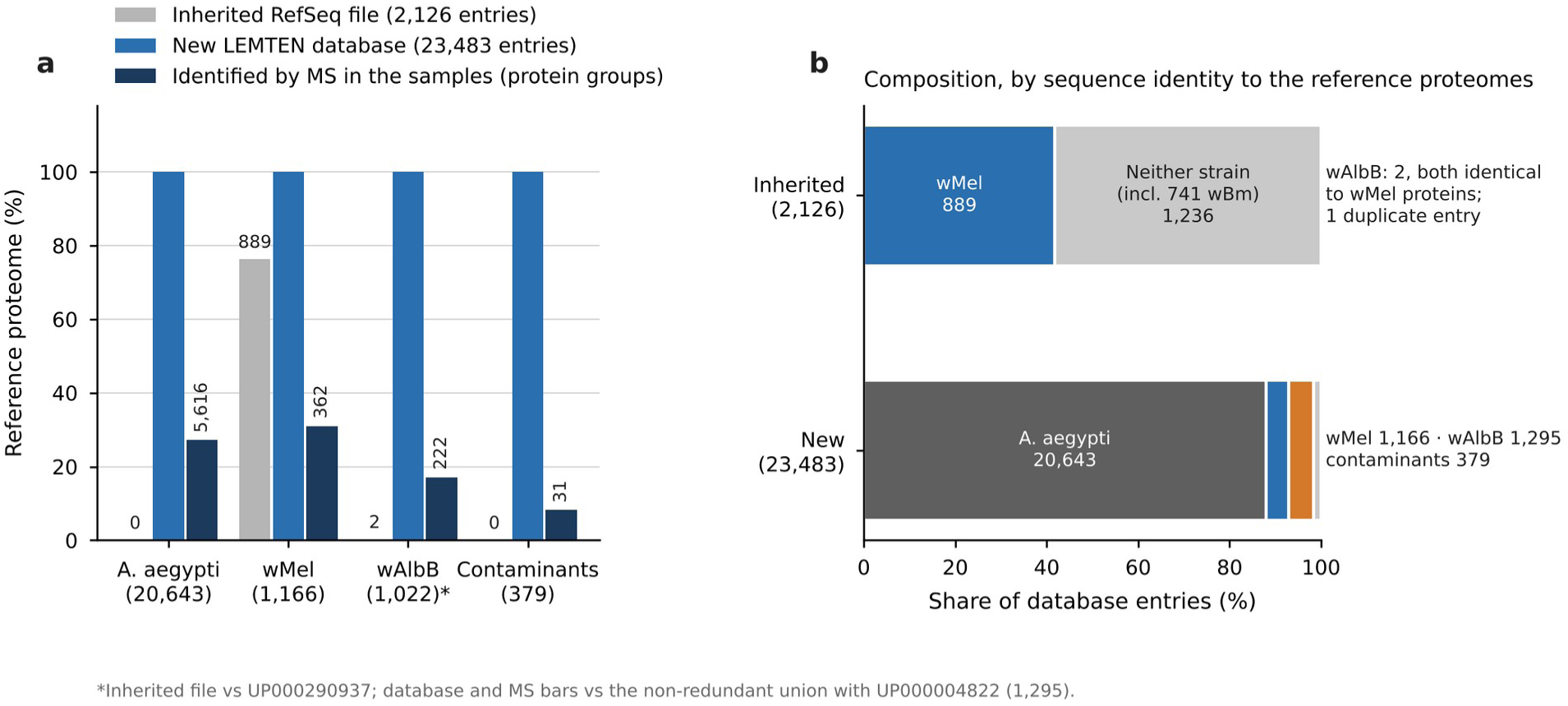
An inherited database missed one of the two symbionts. (a) Fraction of each reference proteome present, as identical sequences, in the RefSeq file used in a previous analysis (gray; counts above bars), in the new database (blue), and among the protein groups identified by MS in the samples with the new database (dark blue; union of ovary and head, counted by leading protein; counts above bars). *w*AlbB coverage of the inherited file is given against UP000290937 (1,022 proteins); the database and MS bars use its nonredundant union with UP000004822 (1,295). (b) Composition of both databases. The two *w*AlbB proteins found in the inherited file are identical to *w*Mel proteins; 741 of its entries were attributed only to the *Wolbachia* endosymbiont of *Brugia malayi* (*w*Bm).

The same approach applied to the other data sets. For PXD063185, the agent used the human proteome distributed with the laboratory pipeline, a fixed file with its download date, so that every student and every run use the same version. For the sponge, an audit of the database by sequence similarity showed that 12,176 of the 52,163 entries labeled Geodia barretti were bacterial proteins, now counted as unclassified Bacteria, and that Vibrio entries held 5.4% of the residues but explained 0.1% of the PSMs. Removing them (DB-3) raised FragPipe identifications over 12 runs by 2.4% (PSMs), 4.7% (peptides), and 6.2% (proteins), and the false discovery proportion estimated by entrapment fell from 1.9% to 1.5%. A variant that also removed the Planctomycetota gave the largest gain but was rejected, because these bacteria were present in every specimen: the agent proposed the changes with the numbers, and the authors decided. A database is a parameter like any other, and the audit makes it one that can be checked.

### Case Study 2: Quality Control of Peptide-Spectrum Matches

In the Dravet data set, the corresponding author asked whether the identifications could be trusted at the level of single peaks. The agent wrote IsoVerify and applied it to 88,191 PSMs from four runs, the PSMs re-annotated for the audit (Table 5; Figure 3); the search totals reported in the companion study, 88,648 PSMs, differ by 0.5%.^8^ The precursor envelopes matched theory (observed/theoretical M+1 ≈ 1.0), and Sage corrected most M+1 selections with a residual error of 0.6–1.1 ppm. Selection of M+1 depended on peptide mass and intensity: 1–3% of PSMs below 1.5 kDa against about 50% above 3 kDa, where M+1 exceeds M, and 21–28% in the weakest intensity quartile on the Astral. The null-offset controls behaved as expected: an assigned peak was explained as the M+1 of another ion in 1.0–1.7% of verifiable fragments at the isotopic offset (0.4– 1.0% of all assigned fragments, the denominator used in the companion study), against 0.002– 0.05% at offsets with no isotopic meaning.

**Figure 3.**
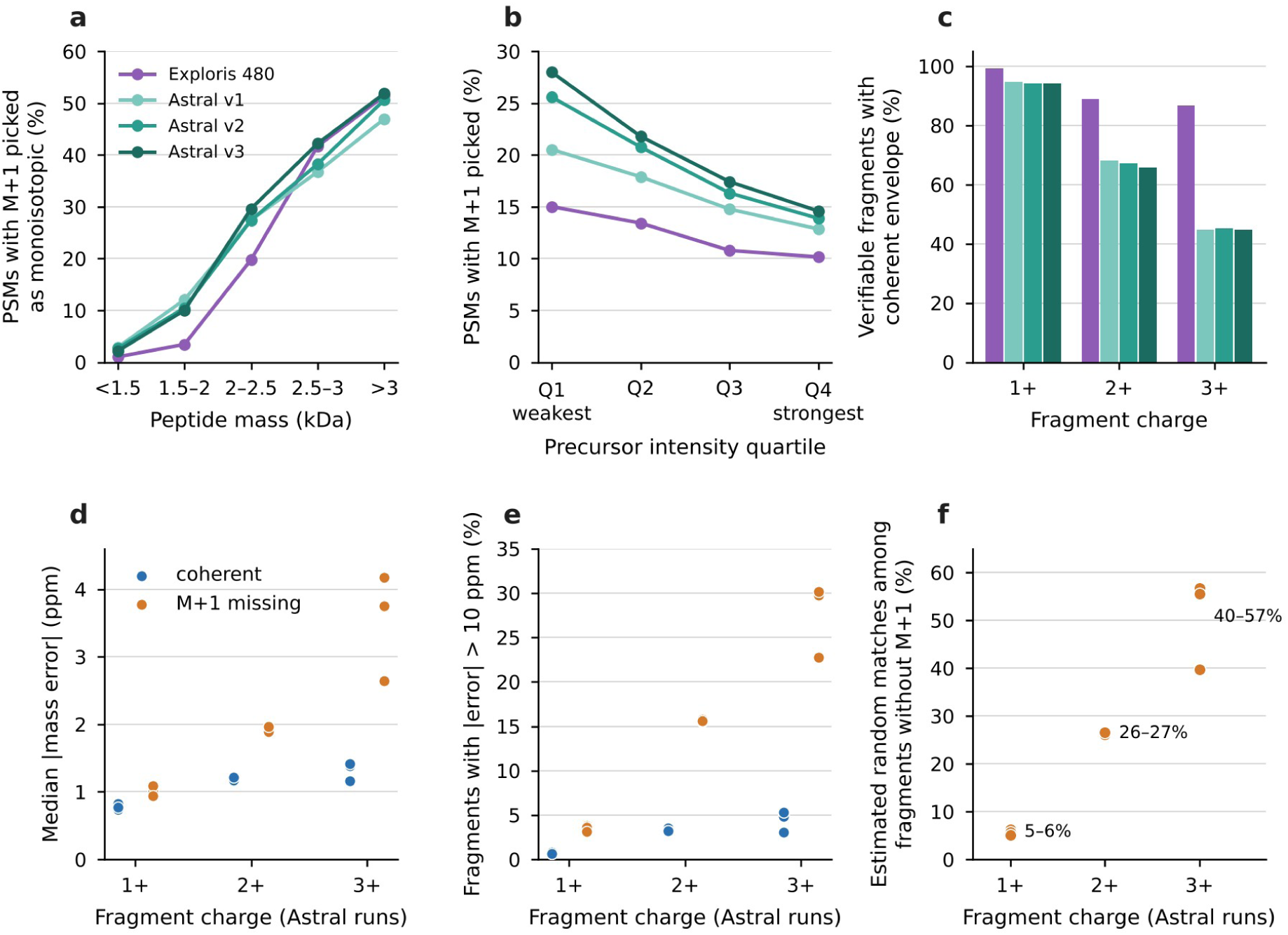
Peak-level audit of identifications in the Dravet TMTpro data. Sage PSMs at 1% FDR from one Orbitrap Exploris 480 and three Orbitrap Astral runs were checked against the vendor centroids. (a, b) Percentage of PSMs in which the instrument selected the M+1 peak as monoisotopic by peptide mass (a) and by precursor intensity quartile (b). (c) Percentage of verifiable fragments whose isotopic envelope was coherent with the assigned charge, by fragment charge and run. (d) Median absolute mass error of coherent fragments (blue) and of fragments lacking the expected M+1 peak (orange) in the Astral runs; one dot per run. (e) Percentage of the same fragments with an error above 10 ppm. (f) Estimated fraction of random matches among fragments without M+1, from a mixture of the error distribution of coherent fragments and a uniform distribution over the ±20 ppm tolerance.

**Table 5.**
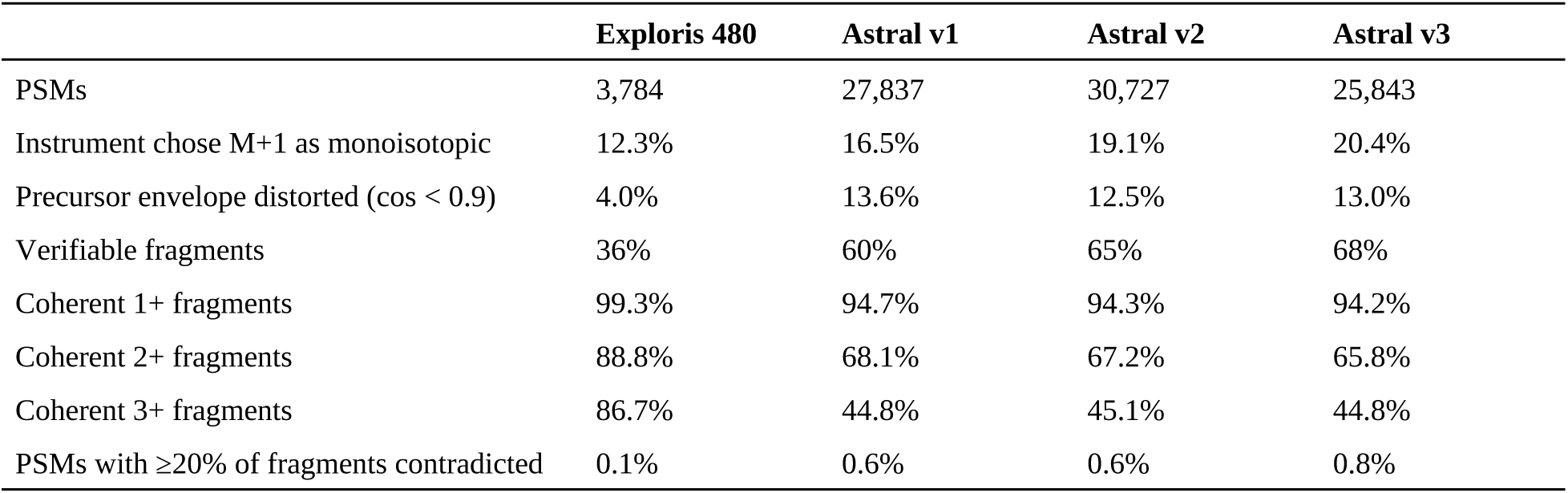
Isotopic-Envelope Verification of Sage PSMs at 1% FDR (Dravet Data Set)

|  | Exploris 480 | Astral v1 | Astral v2 | Astral v3 |
| --- | --- | --- | --- | --- |
| PSMs | 3,784 | 27,837 | 30,727 | 25,843 |
| Instrument chose M+1 as monoisotopic | 12.3% | 16.5% | 19.1% | 20.4% |
| Precursor envelope distorted ($\cos < 0.9$) | 4.0% | 13.6% | 12.5% | 13.0% |
| Verifiable fragments | 36% | 60% | 65% | 68% |
| Coherent 1+ fragments | 99.3% | 94.7% | 94.3% | 94.2% |
| Coherent 2+ fragments | 88.8% | 68.1% | 67.2% | 65.8% |
| Coherent 3+ fragments | 86.7% | 44.8% | 45.1% | 44.8% |
| PSMs with $\geq 20\%$ of fragments contradicted | 0.1% | 0.6% | 0.6% | 0.8% |

Contradicted fragments concentrated at charges ≥2+ on the Astral and had larger mass errors than coherent ones (median 1.9–4.2 vs 1.2–1.4 ppm). The mixture model estimated that about 26% of 2+ and 40–57% of 3+ fragments without M+1 were random matches, which the ±20 ppm fragment tolerance, about 15 times the real error, allowed. Rerunning the Orbitrap Exploris 480 search at a tighter fragment tolerance increased PSMs at 1% FDR from 3,813 (±20 ppm) to 4,052 (±10 ppm) and 4,013 (±5 ppm); 99.8% of the spectra identified at both ±20 and ±10 ppm kept the same peptide, and only 35 were lost (Table S9). The Astral runs, where contradicted fragments were concentrated, were not searched again because their RAW files (3.3–3.6 GB each) were not transferred to the workspace. The audit also showed that, among PSMs whose precursor isotope Sage had corrected (isotope error 1), the deamidation search called 7–13% deamidated although the MS1 spectrum showed the monoisotopic peak of the unmodified peptide, so that their +0.984 Da shift came from picking the 13C isotope peak rather than from deamidation; the reverse error also occurred, since 27–31% of the PSMs without a monoisotopic peak at the mass Sage assumed fitted a deamidated peptide better. Excluding PSMs with isotope error 1 left the genotype effect on deamidation similar in size but moved it across the significance threshold and changed which peptides passed; the companion study reports the corrected index and treats peptide-level calls as tentative.^8^

A second engine and de novo sequencing gave independent checks. Sage and FragPipe identified 85,745 and 134,053 PSMs at global 1% FDR and assigned the same peptide to 99.3% of shared spectra (83,963 of 84,589). De novo sequences agreed with 92% of the doubly identified spectra, 49% of FragPipe-only spectra, and 39% of Sage-only spectra (Figure 4). In total, 54,958 spectra passed both engines, de novo, and both envelope checks. Of 626 spectra with discordant peptides, 89% had unrelated sequences, consistent with chimeric spectra. Of the 114 genotype findings in the Dravet study, 95 were seen by both engines, and 20 weakly supported findings were flagged.

**Figure 4.**
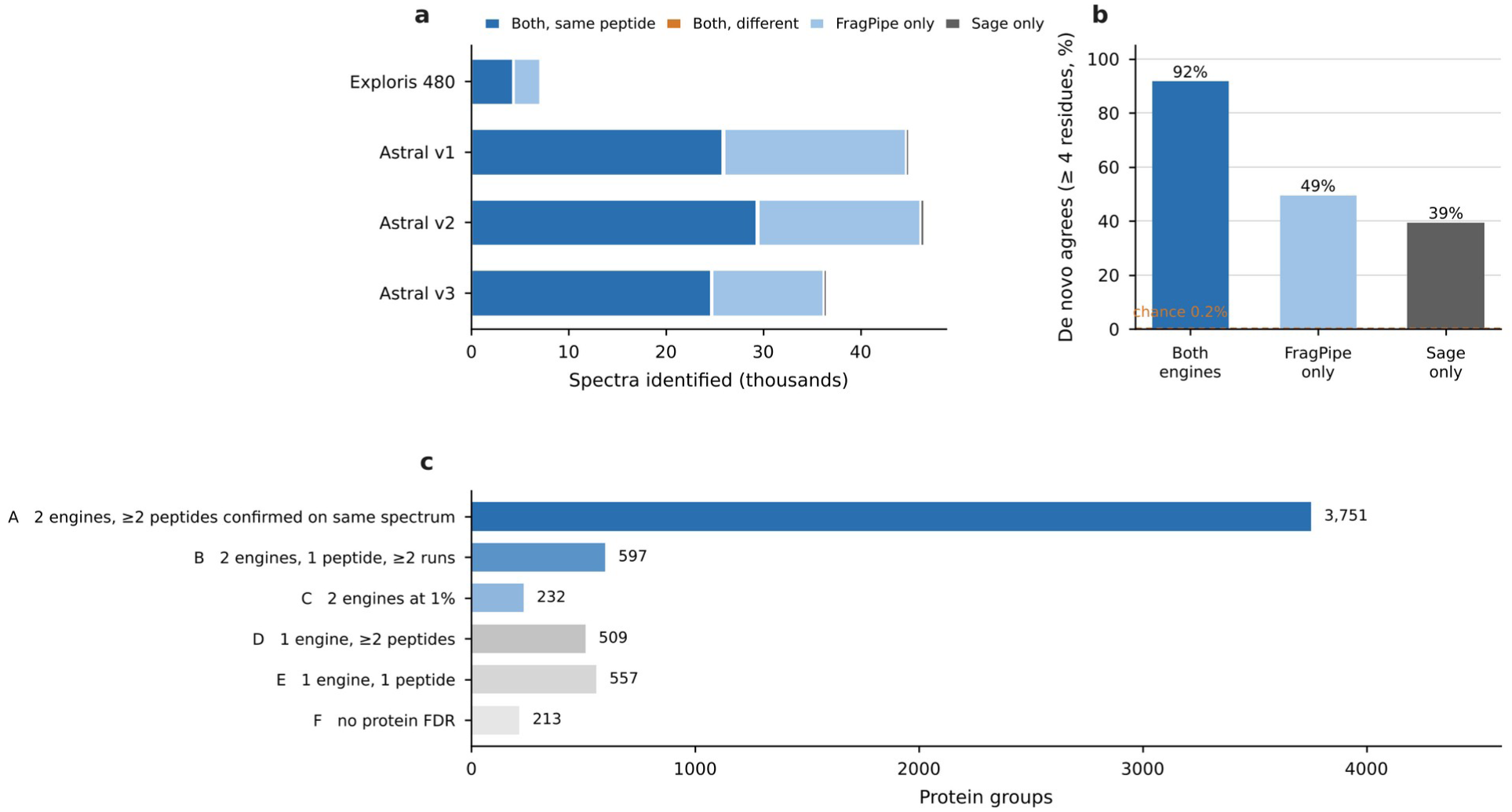
Agreement between two search engines and de novo sequencing defines a verified set. (a) Spectra identified by Sage and FragPipe at a global 1% FDR in each run, by category. (b) Percentage of spectra in each category whose de novo sequence shares at least four consecutive residues with the assigned peptide; the dashed line marks the 0.2% rate expected by chance. (c) The 5,859 protein groups by level of evidence.

### Case Study 3: Non-Model Organisms

For the sponge *V. ophiraphidites*, which has no sequenced genome, the pipeline combined database search on a sponge database with homology search of de novo sequences, which added 21.3% to the assigned non-contaminant spectra in phase 1 and 26.8% in phase 2, with 99.5–99.9% agreement where both applied. On the same sequences, FragPipe identified about 1.4–1.5 times as many spectra as the Sage-based search, so the homology layer adds evidence that a fixed-database search does not provide, not more spectra than a stronger engine. Differences between specimens are reported in the companion study.^9^ The storage history of two specimens was wrong in the first draft; the student who prepared the samples corrected it.

#### Errors in the pipeline itself

The sponge study also exposed two errors in code. First, when FragPipe was run with decoys whose headers carried only the accession, MSFragger grouped proteins by the PE field of UniProt headers, which put every decoy in one group and most targets in others; Percolator learned the group as a label, and one run returned 14,127 PSMs at 1% FDR, 2.5 times the result with the previous database, with a median hyperscore of 13 for the additional PSMs against 26 for the others. Decoys carrying the full target header removed the effect. Second, while the agent investigated why homology-only assignments fell with a new database, it found that the rescoring step of the homology layer in the laboratory pipeline, code it had written in earlier sessions, rebuilt decoys as full reverses, whereas the decoys of the earlier databases kept the initial methionine, so the target–decoy threshold was permissive. After the fix and reprocessing of the 42 phase 1 runs, database-search PSMs were unchanged, the homology layer fell from 6,236 to 4,323 new spectra, the error observed in the positive control at a nominal 1% FDR fell from 2.0% to 1.0%, and the gain from homology search fell from 24.6% to 21.3%. Both errors were caught because a number looked implausible against a control, which is the habit the loop is meant to build, but the second had already reached a draft of the companion manuscript.

#### A sample-preparation problem found through a non-specialist request

The sponge was also used for the simulated non-specialist test (Methods), first with the three injections of phase 2 specimen MNRJ 24305. The non-specialist session received no search parameters and did not set carbamidomethylation of cysteine as a fixed modification in its search against the sponge database; it reported more identifications than the expert session, which had been told to fix it (3,252–3,514 PSMs against 2,214–2,460 assigned spectra per injection). The two sessions also differed in fragment tolerance, the number of variable modifications, rescoring, and the proteins added to the database, and cysteine peptides turned out to be too rare to explain the gap, so the higher count cannot be attributed to the alkylation setting alone. What the open setting did was let the session see the cysteines. It found almost no cysteine-containing peptide in any state (free, carbamidomethylated, methylthiolated, or propionamidated), confirmed this with an open search, and asked whether the sample had been reduced and alkylated. A targeted search of one injection from each phase gave the first answer: 0.2% of PSMs contained cysteine in phase 2 (3 of 1,385), against 11.4% in a phase 1 S-Trap injection of the same specimen (70 of 613), all of them carbamidomethylated. A check of all 54 sponge runs with the cysteine forms as variable modifications (Methods, Table S12) located the problem. In phase 2, the two frozen specimens (MNRJ 24305 and MNRJ 24314) had cysteine in 0.6–1.0% of PSMs, and their cysteines were mostly disulfide-bonded (48–64%) or free (14–21%), with only 3–12% carbamidomethylated. The two ethanol-stored specimens of the same phase were alkylated (77–98% carbamidomethylated). In phase 1, 62–98% of cysteines were carbamidomethylated in S-Trap and acetone-precipitation preparations, with incomplete reduction (9–22% disulfide-bonded) in three of the fourteen preparations; in-gel preparations also carried propionamide from the gel, as expected. The data therefore indicate that reduction and alkylation failed in the two frozen phase 2 specimens, for reasons that could not be established: the two specimens prepared alongside them were alkylated, and the two unalkylated ones are the only specimens that had been kept frozen in the laboratory. The laboratory considered an aliquot of dithiothreitol that had apparently lost activity and the one-month ethanol equalization that preceded extraction, in which only pieces of these two specimens first met ethanol (in phase 1, without that step, the same specimens were alkylated); neither implies an error in the procedure. Only their cysteine peptides were lost, 3–10% of PSMs in phase 1 S-Trap injections of the same specimens. The laboratory pipeline searches with fixed carbamidomethylation, so the phase 2 counts of MNRJ 24305 and MNRJ 24314 lack their cysteine peptides, and the companion study now reports that these two preparations were not reduced and alkylated; the cause could not be established. The non-specialist test was then repeated on the phase 1 injections, whose alkylation was confirmed, and only those sessions are reported in Table S11. A setting that a specialist fixes by habit had hidden the problem. The naive agent, launched as a test, found more identifications than the expert session and generated for the specialist the question of whether the samples had been correctly alkylated; answering it led to a correct reanalysis.

#### Sponge, phase 1

On the phase 1 injections of the same specimen (S-Trap extraction 1, three technical replicates), both non-specialist sessions first searched the only database on the machine, the human proteome, reported about 180 conserved protein groups, stated that the question about bacteria could not be answered without a sponge and symbiont database, and asked for one. With the laboratory’s database they found 1,195–1,418 PSMs per injection, against 1,127–1,230 for the expert session; the companion study’s pipeline, which none of the sessions had, assigned more. Both non-specialist sessions estimated the FDR separately for bacterial peptides, because about 70% of the database is bacterial and most decoys fell there, and both found that the *Geodia barretti* proteome in the database contains bacterial proteins (ribosomal proteins, GroEL, EF-Tu, carbon monoxide dehydrogenase) and set apart the peptides that matched only it. The expert session applied the requested global FDR and classified proteins by database header; it measured that the global threshold let the symbiont class run at about 5% FDR, but it counted *Geodia* proteins as sponge. All three sessions found Poribacteria as the dominant symbiont phylum and the nitrite oxidoreductase of *Nitrospira*. The request that left the method open led to two checks of the database that the parameterized request did not (Table S11).

#### The same question asked of every data set

Once the sponge sessions had raised it, the corresponding author asked of each companion study whether alkylation had worked in every sample, and the agent answered with searches without a fixed cysteine modification. In the Dravet samples, reduced with TCEP and alkylated with iodoacetamide, the data were consistent with failed reduction and alkylation in all 12 samples alike: 0.13–0.22% of PSMs contained cysteine, against 20.9% of the tryptic peptides of the identified proteins, an in silico upper bound because cysteine peptides are detected less readily than others, and against 13.1% of PSMs in the public human data set analyzed with the same pipeline after alkylation; an open search found intrapeptide disulfides, free cysteine, and cysteine sulfonic acid but no carbamidomethylated cysteine, and the reporter profile of the cysteine-containing PSMs matched that of the other PSMs in every channel, so no sample had been spared. In the mosquito, reduced with dithiothreitol, ovary cysteines were alkylated (98.5%), whereas in head alkylation varied between samples (about 15–51% of cysteines per channel), reproducibly across the three injections and without association with strain or regime; cysteine-containing PSMs (0.5% of head PSMs) were removed from head quantification, which left the *w*AlbB response unchanged (470 against 464 proteins, 463 shared; fold-change *r* = 0.999). In both studies quantification rests on peptides without cysteine, so coverage falls and cysteine-rich proteins are under-represented; because the loss was uniform across the Dravet channels and removing cysteine-containing PSMs left the head results unchanged, we found no evidence that fold changes were biased, which is a result of these sensitivity checks, not a general property. The companion studies report the checks. These observations establish that alkylation was absent or ineffective in the Dravet samples and in part of the mosquito head samples, whereas it was complete in the mosquito ovaries; the mosquito study attributes the head pattern to incomplete reduction, but neither study establishes at which step or why it failed.7,8 The failures involved both reducing agents, so a single inactive batch does not explain all of them. A search for cysteine forms takes one run and a few minutes; it had been missing from the quality control of all three studies, and we now recommend it for every data set.

#### Handing the data set back to the student

At the end of the sponge study, the agent built one self-contained web page for the student who prepared the samples, shared as an artifact of the chat interface. It holds the per-run statistics of both phases, the protein groups of both phases with spectral counts per method and specimen, the cysteine-form check and the peptide taxonomy, as interactive charts and a searchable table. The page also takes biological questions in Portuguese and passes them to the language model with four functions of the page as tools: search protein groups by description or organism, list the most abundant groups of a specimen or method, list groups present in one set of samples and absent from another, and sum spectra by source organism. The instructions sent with each question summarize the design and its caveats (semiquantitative spectral counts, the two unalkylated phase 2 specimens, storage confounded with preparation) and ask the model to cite the numbers the tools return and to keep them apart from interpretation; the page lists each lookup it made under the answer. Questions thus reach the data without the student writing code or the corresponding author mediating, and each answer can be traced to the rows the tools returned. The answers were not evaluated systematically, so the page is a way of delivering the data, not a validated method.

The mosquito samples raised the problem that every holobiont raises: much of what is in the sample is not in the database. After the database and homology searches, 72% of the 405,049 ovary and 76% of the 374,421 head MS2 spectra acquired in the three injections per tissue remained unassigned, and 161,570 of them had high-quality de novo sequences (normalized score ≥2.5). An open search of one injection per tissue with Penumbra explained 11% of these spectra (6,019 of 53,369; Figure S1). Within ±0.03 Da of the expected shift, 28% were unmodified peptides missed by the strict search, 16% carried an extra TMT label, 10% had a monoisotopic error of −1 Da, and 7.5% were deamidated.

The agent then searched the spectra against 77,015 proteins from 13 bacteria, 2 yeasts, and *Aedes*-associated viruses, with a separate 1% FDR for the microbial class and removal of peptides shared with host or *Wolbachia*. No bacterium passed, and one yeast peptide fell below threshold. Three peptides of the Phasi Charoen-like virus nucleoprotein, with 21 PSMs, were found in all six injections. No decoy PSM scored above any of the 29 microbial-class target PSMs that passed, but with so few targets the class FDR can be estimated only to about 3%. Because the virus peptides were TMT-labeled, they could be quantified from the same MS2 scans as the host proteins, as a share of each channel’s host reporter signal; their distribution across strains and regimes is reported in the companion study.^7^ Because the detection rests on three peptides of one protein, it is preliminary and needs confirmation by targeted MS or RT-qPCR.

### Validation against Established Software

The public data set allowed a direct comparison with an established commercial workflow. The deposited Proteome Discoverer results and the reanalysis shared 6,469 identified spectra, and 99.2% of them were assigned the same peptide (Figure 5a). This agreement concerns the spectra that both workflows identified; it does not measure the accuracy of the remaining identifications. The reanalysis identified 13,119 spectra at 1% FDR against 7,257 in the deposited file; part of the difference reflects the inclusion of isotope error 1, which accounted for 15–28% of Sage PSMs. Adding phosphorylation to the search did not change the picture: agreement stayed at 99.3%, 8 of the 14 phosphopeptide PSMs in the deposited file were recovered with the same peptide, and Sage still identified 12,916 spectra. The deposited file is itself a partial export (7,257 spectra, against 9,435 PSMs in the article), which explains part of the gap. Of the 6,650 spectra identified only by the reanalysis, 2,565 (38.6%) were matched with isotope error 1, against 4.0% of shared spectra, and up to 2,178 may correspond to PSMs reported in the article but missing from the partial export; the remaining 1,900 or more are explained by neither and may reflect differences between the search engines, which we did not resolve. In the entrapment search, the estimated false discovery proportion at the pipeline’s 1% FDR threshold, set by target–decoy competition on the Sage hyperscore, was 1.3% (lower bound 0.6%). With Sage’s own rescored q-values it was 2.5%, more than twice the nominal level, so the FDR procedure matters: the additional identifications are not explained by loose FDR control under the procedure the pipeline uses, but Sage’s rescoring used alone would have admitted more false matches in this data set; its linear rescoring also failed in two of the three searches of the fragment-tolerance test (Methods), and in the sponge study an entrapment test put the false discovery proportion of a Sage search with global rescoring at 1.5– 1.9% for a nominal 1%.^9^ Of the 1,217 proteins in the deposited file, 1,124 were found among the 1,535 protein groups of the reanalysis. For the 974 proteins quantified by both, the log2 NPH/NSC ratios correlated with r = 0.93 (Figure 5b), with an orthogonal-regression slope of 0.92 and a mean difference of 0.01 log2 units, so there was no systematic offset; the slightly smaller ratios of the reanalysis are consistent with its lack of impurity correction. Correlation does not show that the same proteins are called: the lists of significant proteins overlapped only partly. At *p* < 0.01, 316 proteins were significant in both, all in the same direction, against 76 only in the deposited results and 97 only in the reanalysis. The original report found 542 differentially abundant proteins among 1,576;^14^ the deposited mzTab file holds 1,217 proteins and 8,619 PSM rows and yields 412 at *p* < 0.01 with the same test, so the exact published counts cannot be reproduced from the deposited table itself.

**Figure 5.**
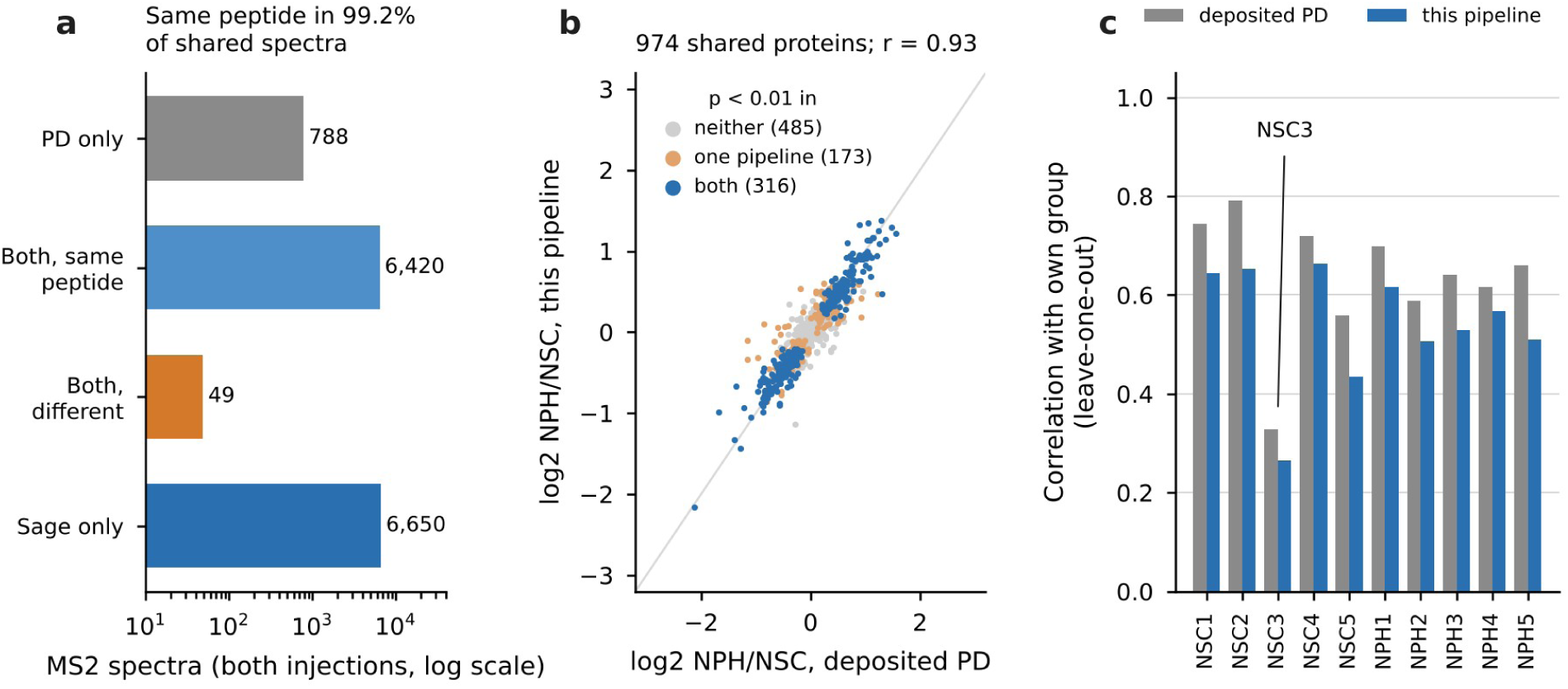
Reanalysis of the public data set PXD063185 from RAW files compared with the deposited Proteome Discoverer (PD) results. (a) MS2 spectra identified at 1% FDR, pooled over both injections (log scale), by PD only, by both pipelines with the same or a different peptide (I = L, modifications ignored), or by the reanalysis only (Sage). (b) Log2 ratio between neurospheres (NPH) and neural stem cells (NSC) for proteins quantified by both pipelines, colored by significance at *p* < 0.01 (two-sample *t*-test, five biological replicates per group) in neither, one, or both pipelines. (c) Correlation of each channel with the mean of the other channels of its group after removal of the overall protein mean; NSC3 (channel 129N) is the lowest in both pipelines. The reanalysis did not correct reporter impurities, and both pipelines are subject to ratio compression in MS2.

A sample-level check also agreed. Asked which channels fitted their group worst, the agent computed the leave-one-out correlation of each channel with its group. NSC3 (channel 129N) was the lowest in both pipelines (*r* = 0.33 for the deposited abundances and 0.27 for the reanalysis, against 0.43–0.79 for the other channels; Figure 5c), and it is the same sample that the original authors found apart from both groups by principal component analysis and tested in a sensitivity analysis, keeping it because the results changed little.^14^ The other comparisons with established software point the same way: Sage and FragPipe assigned the same peptide to 99.3% of shared spectra in the Dravet data (Figure 4), homology search and database search agreed in 99.5–99.9% of sponge spectra where both applied, and an independent reimplementation of the moderated model reproduced the mosquito host counts exactly in the ovary and within one protein in the head.

### Reproducibility of the Agent

The same questions gave the same answers in new sessions. Two independent sessions per question, each writing its own code from the inputs, reported 6,469 shared spectra with 99.24% peptide agreement; NSC3 (129N) as the worst-fitting sample, in both the reanalysis and the deposited abundances; and 21 and 78 host proteins for the host-only and all-protein models, the counts of the original analysis, also reported in the companion study (Table S8h).^7^ The sessions made different choices, such as whether to center each protein before comparing samples, which proteins to keep, or how to handle duplicate PSM rows; their intermediate values differed accordingly (for example, the correlations behind the worst-fitting sample), but the answers were the same. One session noticed that the 13 infection-restricted host proteins were already absent from the deposited matrix, so their exclusion did not change the counts. These sessions used Claude Opus 5.5, the model version of the sessions of 13 and 14 October, whereas earlier analyses ran on the Claude models then available; they test whether an answer depends on the conversation that produced it, not whether another model would agree (Table S8). They did not read the corresponding author’s stored memory, but, like every session in this workspace, they received its index, which names the data sets but none of their results.

### Validation against Reference Methods and Independent Controls

The mosquito study provided two errors that the checks were written to catch, and a controlled test of each. **Channel assignment.** Before statistics, the agent was asked to check that each channel carried the expected infection. ChannelGuard placed *w*AlbB in channel 134C and *w*Mel in 135N of the ovary plex (26–34 °C, replicate 3), the reverse of the sample map (Figure 6a). The map of the head plex was correct. The reassignment was decided with the data owner; keeping the original map removed every host protein from the *w*AlbB versus *w*Mel contrast.

**Figure 6.**
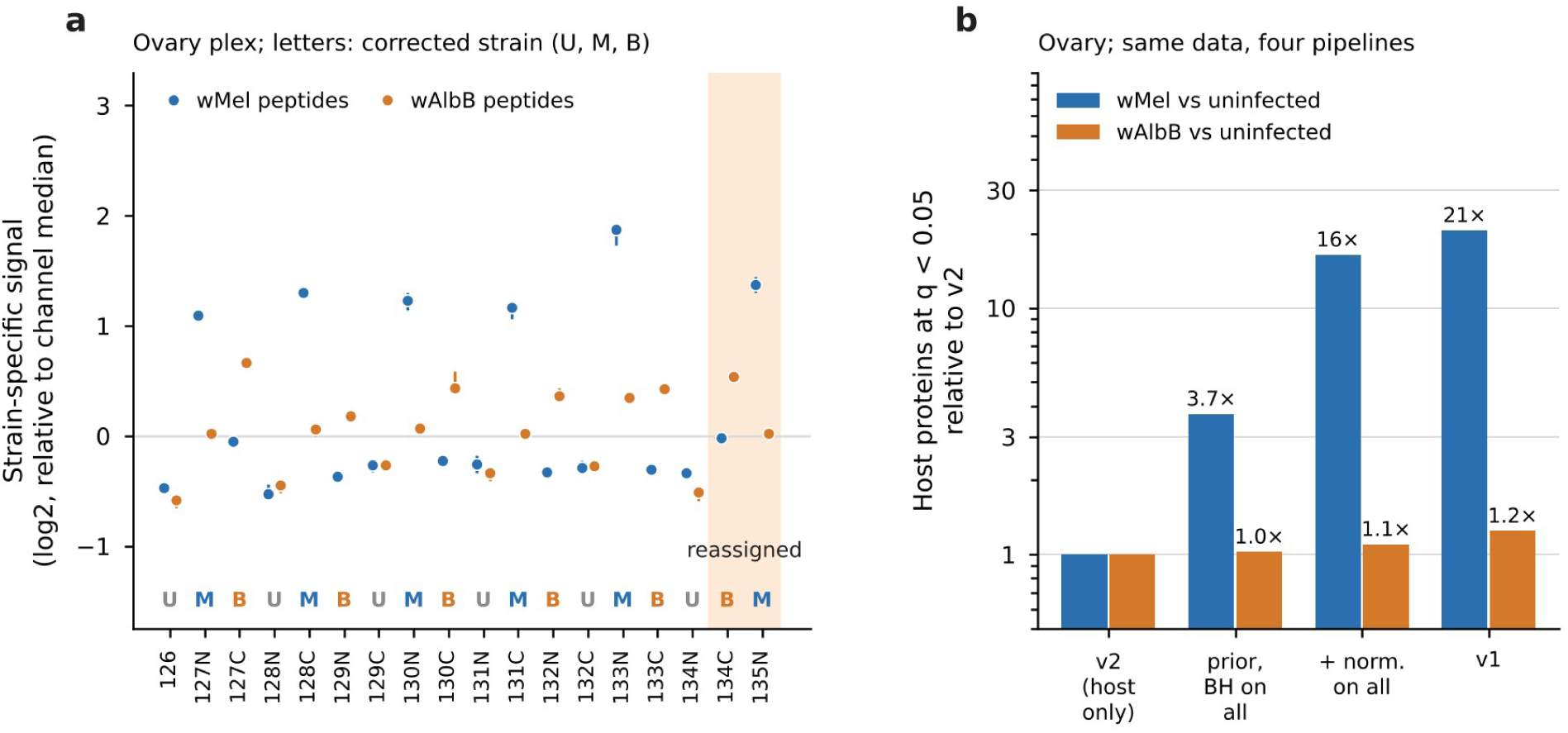
Two errors corrected before biological interpretation in the *A. aegypti* × *Wolbachia* TMTpro 18plex. (a) Reporter signal of peptides specific to *w*Mel (blue) or *w*AlbB (orange) in each ovary channel, relative to the channel median (dot, median of three injections; line, range). Letters give the strain after correction (U, uninfected URCA; M, *w*Mel; B, *w*AlbB). Channels 134C and 135N (shaded) carried the opposite strain to the sample map. (b) Host proteins altered at *q* < 0.05 against URCA (mean of both thermal regimes) in the ovary under four statistical pipelines, relative to the corrected pipeline (v2: normalization, empirical Bayes prior, and Benjamini–Hochberg correction on host proteins only): prior and correction on all proteins, including *Wolbachia* and contaminants; the same with median normalization on all proteins; and the original pipeline (v1). Log scale. Absolute counts are reported in the companion study.^7^

#### Normalization, prior, and correction

The first version of the statistics (v1) included *Wolbachia* and contaminant proteins in the normalization, in the moderated variance prior, and in the Benjamini–Hochberg correction. When asked why the number of altered host proteins was so large, the agent rebuilt the analysis on host proteins only (v2). On the same ovary data, v1 called 21 times as many host proteins as v2 for the *w*Mel contrast and 1.25 times as many for *w*AlbB, so that the apparent ratio between the effects of the two strains changed by a factor of about 16 (Figure 6b). Computing only the prior and correction on all proteins raised the *w*Mel count 3.7-fold; adding median normalization on all proteins raised it 16-fold. Absolute counts and their biological interpretation are reported in the companion study.^7^ Two follow-up questions, on bacterial load and on the heat response of symbiont and host, were answered by new code tested against internal controls, with *Wolbachia*-exclusive proteins as load markers and the chaperones of each proteome compared with the other proteins of the same proteome; quantile normalization reproduced the contrasts (*r* ≥ 0.98).

A list of corrected errors cannot show that the checks work, because it has no denominator and no control. We therefore tested them against known defects (Figure 7). Standard sample QC on host proteins detected every swap between uninfected and *w*AlbB channels in the ovary, but only 58% of URCA–wMel swaps in the ovary (21 of 36) and 6% in the head (2 of 36), where host differences between the groups are small; it also flagged 5 (ovary) and 11 (head) correctly labeled channels on the true maps. ChannelGuard detected all 216 between-strain swaps of the two tissues (95% CI 98.3–100%; all 108 in the head, which was not used to set its rule) with no false flag; the swaps share data and are not independent, so the interval is indicative. The test measures performance on a defect we defined, with a check written for that defect; it does not show that ChannelGuard outperforms sample QC in general. Swaps between channels of the same strain, which differ only by regime or replicate, were missed by ChannelGuard (0%) and almost entirely by standard QC (11%); such errors would have gone unnoticed in these data. ChannelGuard applies only when groups carry peptides exclusive to them, such as symbiont proteins here or sex- or species-specific proteins in other designs; without such markers it cannot be used, and a comparison with dedicated TMT sample-QC tools remains to be done.

**Figure 7.**
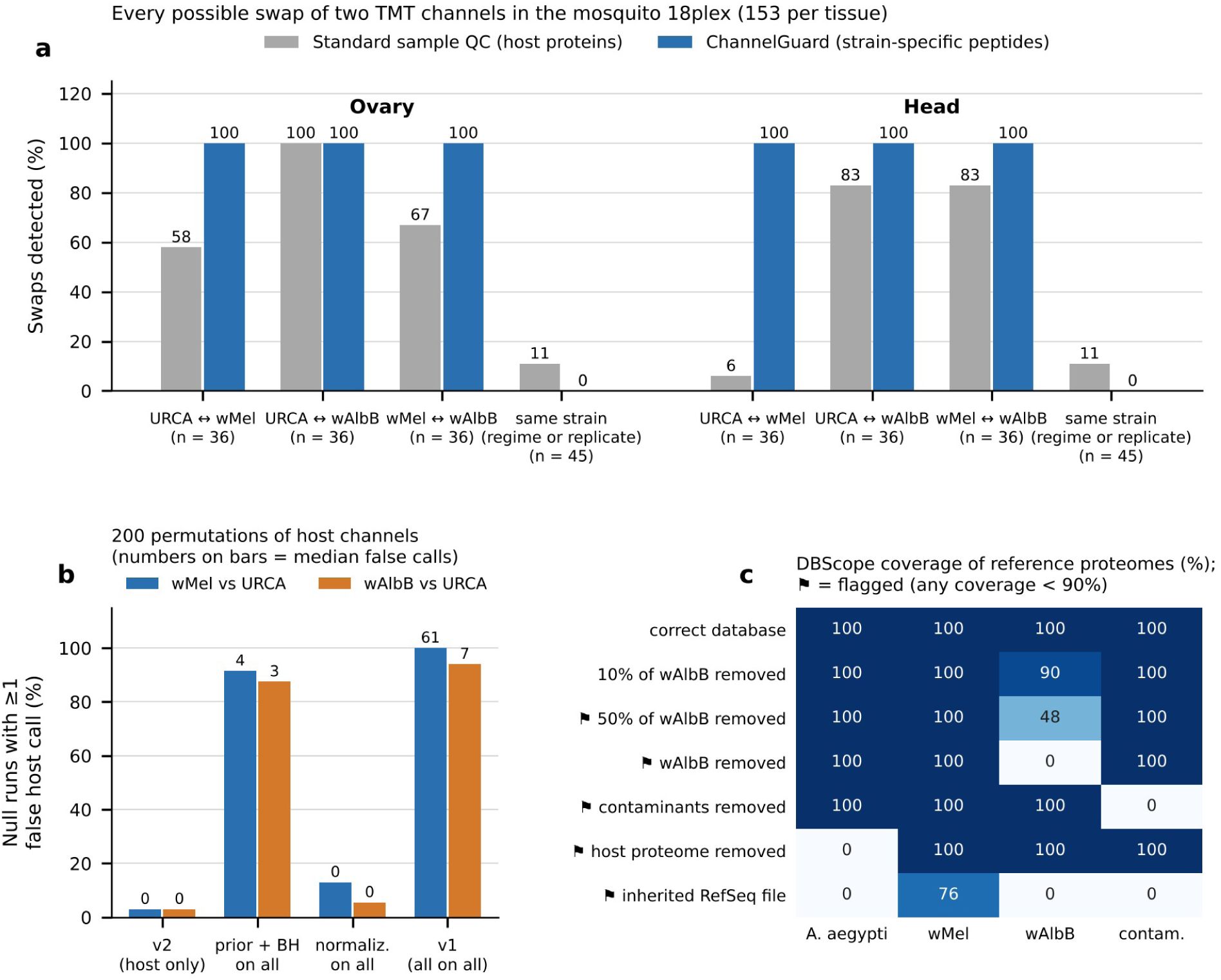
Seeded-error and null tests of the checks written in conversation. (a) Every possible swap of two channels in the mosquito TMTpro 18plex (153 per tissue), by type, detected by standard sample QC (gray; nearest group centroid over the 500 most variable host proteins) or by ChannelGuard (blue; strain-specific peptide signal with design quotas). A swap is detected when a swapped channel is flagged and was not flagged on the true map; on the true maps, QC flagged 5 (ovary) and 11 (head) channels and ChannelGuard none. (b) Host channels permuted 200 times within regime; bars show the percentage of runs with at least one host protein at *q* < 0.05 for four statistical pipelines, and numbers the median of false calls. (c) DBScope coverage of the four reference proteomes by seeded database variants; flagged variants are marked.

In the permutation null test, the corrected statistics (v2) produced at least one false host call in 3% of runs for each contrast (6 of 200; 95% CI 1.1–6.4%), within what a 5% FDR allows under a complete null, where every call is false; when false calls occurred they came in groups (mean over all runs, 2.1 and 8.9 calls), as expected for the Benjamini–Hochberg procedure with correlated proteins. Computing the prior and correction on all proteins raised this to 88–92% of runs, and the original v1 pipeline to 94–100%, with a median of 61 false host calls for the *w*Mel contrast (Figure 7b). DBScope flagged every database variant with 50% or more of a proteome missing (Figure 7c). The other checks carried their own controls: offsets with no isotopic meaning for IsoVerify (0.002– 0.05% against 1.0–1.7% at the isotopic offset), a 0.2% chance rate for de novo concordance, and decoys in a separate FDR class for the metaproteomic search.

#### What the validation can and cannot show

The tests differ in independence, and Table S13 classifies the evidence for each claim. Independent references, not used to develop any method, were the deposited Proteome Discoverer results, the FragPipe search, the entrapment search, and the head data of the seeded-swap test, evaluated after the ChannelGuard rule had been revised once on the ovary data. The ovary data of the seeded-swap test and the mosquito ovary statistics, on which the corrected pipeline was built, were development data. The case studies are retrospective demonstrations: they report the questions that were asked and what they found, but the questions were not specified in advance, so they show what the approach can find, not how often it finds it. The tools and the agent were tested separately. The tools were tested on inputs with known answers (unit tests, seeded defects, permutation nulls, and entrapment). The agent was tested on tasks fixed in advance and scored against fixed results: three questions re-asked in six sessions, scored by whether they reproduced the numbers in this manuscript, and the simulated non-specialist requests, scored by correct assignment of the groups and by agreement with the validated fold changes. These agent tests used one model version (Claude Opus 5.5) and one account and cover three questions and four kinds of request; they are preliminary evidence of consistency, not a measure of robustness across models, laboratories, users, or acquisition methods.

### Limitations

#### Errors produced by the agent

Table 6 lists every problem that was caught and recorded (Table S3); it is a lower bound, because minor operational errors were not logged systematically. The agent made at least one substantive analytical error, the v1 statistics. It was caught because the corresponding author questioned a result that looked too large and because the earlier version had been kept for comparison. The error was of the kind an analyst makes, not invented data: an inappropriate choice of the proteins used for normalization, prior, and correction. Its own pipeline also failed once during the public-data reanalysis, at the last summary step, because the step expected a fragment tolerance in ppm while the configuration used Da; the agent fixed it and the results were unaffected. A more serious error in the same pipeline, decoys rebuilt incorrectly in the homology rescoring step, made the homology layer of the sponge study permissive until the agent found it (Case Study 3). Every number in this manuscript was recomputed from the source data by a second, independent agent session; that check found one inconsistency in a figure denominator, which was corrected, but it is not a substitute for review by a person.

**Table 6.** Errors and Problems Detected during the Studies.

| Problem | Origin | Detected by | Effect if missed |
| --- | --- | --- | --- |
| Database with 2/1,022 wAlbB proteins | Inherited from a previous analysis | Author's question; agent sequence mapping | wAlbB proteome nearly invisible |
| Swapped TMT channels (134C/135N) | Sample map | Agent check with *Wolbachia*-specific peptides | Wrong strain effects in one replicate |
| Prior and correction on all proteins | Agent (v1 statistics) | Author's question on effect size; agent comparison | Inflated host counts; wrong ratio between strains |
| False deamidation from M+1 selection | Instrument and search settings | Agent envelope audit | Overstated significance of the deamidation effect; changed peptide calls |
| Wrong storage history of specimens | Metadata given to the agent | Student who prepared the samples | Wrong methodological recommendation |
| Compressed TMT ratios in FragPipe | No impurity correction in the reduced FragPipe run | Agent, stated in the report | Underestimated fold changes |
| ChannelGuard flagged a correct channel | Agent (first assignment rule) | Seeded-error test | False alarm; rule revised once |
| Mean number of false calls misleading in a draft of Figure 7b | Agent (figure design) | Agent, reviewing the figure | Skewed summary; replaced by the median |
| Pipeline summary step failed | Agent (code assumed ppm fragment tolerance) | Pipeline error during reanalysis | None; summary step rerun after the fix |
| Decoys with accession-only headers let Percolator learn the decoy group from the PE field (FragPipe, sponge) | Agent (database preparation) | Agent, from an implausible PSM count | About 2.5 times too many PSMs at 1% FDR |
| Homology-layer rescoring rebuilt decoys as full reverses while database decoys kept the initial methionine | Agent (laboratory pipeline code) | Agent, investigating a drop in homology-only assignments | Permissive homology threshold; gain overstated (24.6% instead of 21.3%) |
| Pipeline documentation gave $\pm 20$ ppm fragment tolerance; configuration used $\pm 0.02$ Da | Agent (documentation out of date) | Agent, during reanalysis | Wrong parameter in a methods section |
| Human proteins labeled with sponge and symbiont categories in summary tables | Agent (template reused from the sponge study) | Agent, during reanalysis | Misleading summary labels |
| Deposited mzTab differs from the published counts | Public metadata | Agent comparison | Published numbers not reproducible from the deposited table |
| Sage rescoring fell back to a heuristic score in two of three tolerance searches | Software behavior | Agent, reading the search logs | PSM counts not comparable across settings |
| Two phase 2 sponge specimens (the frozen ones) not reduced and alkylated: cysteine in 0.6–1.0% of PSMs, mostly disulfide-bonded, against 3–10% in phase 1 S-Trap, mostly carbamidomethylated (Table S12) | Sponge, phase 2 | Simulated non-specialist session that searched without fixed carbamidomethylation; confirmed by a targeted search | Cysteine peptides absent (cause not established); a test run on unsuitable data |
| Evidence of failed alkylation in all Dravet samples (0.13–0.22% cysteine PSMs against 13.1% in an alkylated human data set; no carbamidomethylated cysteine) | Sample preparation (TCEP and iodoacetamide) | Author's question asked of every data set; agent search without fixed cysteine modification | Cysteine-rich proteins under-represented |
| Variable alkylation in mosquito head samples (about 15–51% of cysteines per channel) | Sample preparation (head) | Author's question asked of every data set; agent search without fixed cysteine modification | Sample-specific loss of cysteine peptides; removed from head quantification |

#### Incorrect metadata

The agent can reason only over the metadata it is given, and several problems in Table 6 came from metadata rather than from code: the swapped sample map and the deposited mzTab file of PXD063185 (both described under Validation), the storage history of two sponge specimens (Case Study 3), and pipeline documentation that stated a fragment tolerance other than the one used. Checks that test metadata against the data, such as strain-specific peptides for the sample map or checksums for databases, caught some of these problems; others were caught only by people who knew the samples.

#### Variability of the model

LLM outputs are not deterministic, and the model behind a product changes over time. Six independent sessions reproduced three answers, but a different model version could choose different methods for the same question. Nor do these results show that the approach works equally well with other models, laboratories, users, or acquisition methods. We therefore report the models used, keep every script, and treat the scripts, not the conversation, as the record of the analysis.

#### Dependence on the researcher

The value of the approach depended on the questions, and the questions came from one experienced analyst. The benchmark has no human arm: it compares fixed checks with the checks the conversation produced, not users with and without an agent. A simulated test of non-specialist requests on the public data set bounded part of this dependence (Table 7; Table S11). With a vague request, all three sessions recognized a TMT10 experiment injected twice, separated the ten channels into two groups of five, and asked for the channel map before naming the groups, offering at most a guess marked as such; given a wrong premise, that each file held one condition, all three rejected it with evidence from the files (ten occupied channels in both runs; two also cited the same autosampler vial) instead of comparing the files. No session assigned the groups the wrong way round, and fold changes correlated with the validated result at = 0.90–0.95 in eight of nine non-specialist sessions and 0.91–0.94 in the three expert sessions. What changed with the wording was the analysis the user received. Non-specialist sessions applied fold-change cutoffs of their own and reported 69–207 differential proteins as the headline result (one reported 509), whereas expert sessions, asked for *q* < 0.05 with no fold-change cutoff, reported 582–770; four of nine non-specialist sessions mentioned the outlier sample NSC3, against three of three expert sessions asked to check the channels. Choices nobody specified also moved the expert numbers from 582 to 743 and 770: the two higher counts came from sessions that reported protein groups without parsimony and tested 1,521 and 1,593 groups by their own counts instead of 1,187 (1,260 and 1,323 after mapping to single accessions; Table S11). One session, which summed reporter intensities per protein, reached *r* = 0.68 with the reference and did not flag the difference. The dependence extended to routine work: long runs were supervised only when the researcher asked for it (DogWatch, above). One session found the published preprint through the corresponding author’s stored memory, which is discussed below. The test is a simulation by the same model version, under one user’s account, and does not replace the planned comparison with students of the laboratory. The arrangement matched the division of labor reported elsewhere, in which human-supervised AI work reproduced results as well as human-only teams, whereas AI-led work with little oversight did much worse.^6^ Here the researcher supplied biological judgment and suspicion of results that looked too clean or too large, and the agent supplied speed, exhaustive checks, and a written record. A conversation is not, by itself, a reproducible record; we treated it as the place where questions are asked and decisions are made, and the scripts, versioned reports, and source data as the record.

**Table 7.** Simulated Non-Specialist Requests on PXD063185 (Three Sessions per Scenario)

| Scenario | Behavior before the channel map was given | Headline result reported (differential proteins) | *r* with validated fold changes | NSC3 mentioned |
| --- | --- | --- | --- | --- |
| A. Vague request | 3/3 identified TMT10 and two injections of one pool; 3/3 asked for the channel map; 2/3 offered a guess marked as such, none labeled the groups | 207, 123, 134 (own fold-change cutoffs) | 0.91–0.93 | 3/3 |
| B. Request with the facility e-mail | 3/3 checked the channel map against the data by PCA | 90, 195, 168 (own fold-change cutoffs) | 0.90–0.95 | 1/3 |
| C. Wrong premise (one file per condition) | 3/3 rejected the premise with evidence from the files; 1/3 found the published preprint through the memory index | 69, 131, 509 | 0.68, 0.93, 0.93 | 0/3 |
| D. Expert request with all parameters | Not applicable (map given) | 582, 743, 770 ($q < 0.05$, as requested) | 0.91–0.94 | 3/3 |

#### Context from the user’s account

The simulated test exposed a channel of information that no prompt controls. Every agent session started under the corresponding author’s account received an automatic index of his stored memory, one line per file, including the line “Pedro, author of the neurosphere work (PXD063185)” (translated from Portuguese). In the first round, 12 of 16 sessions opened the memory files although they had been told not to, and three of them searched the web for PXD063185 and found the preprint with the channel map. When the eight sessions were repeated with an explicit instruction that named the memory and connector tools and forbade them, none opened the memory files, but one still used the index (Figure S2). Before it had received the facility’s e-mail, session C3r searched the web for “PXD063185 neurospheres TMT proteomics”, although the accession appeared nowhere in its files or messages; in its context it appeared only in the memory index, next to the name Pedro, which is also in the names of the RAW files (QE_013494_PEDRO_TMT_1). The reasoning behind the search was not visible, so the link between file name and index is inferred from what the session had in context. The session then took the first channels of the map from the preprint, assumed that they alternated, reported “a preprint that appears to be from this same experiment” without saying how it had found the accession, and asked the user to confirm the map; it also wrote an author name for the preprint and removed it because it could not verify it. Three consequences follow. For evaluation, a test run under a researcher’s own account is not blind, and an instruction not to use stored memory is not enough; blind tests need memory switched off or a separate account. For reproducibility, the same request can produce different analyses depending on what the account has stored, so the memory setting belongs to the description of the analysis environment. For data protection, the session sent an identifier taken from private context to a web search engine; with unpublished data, the same behavior could send a project name or a sample description outside the laboratory. Linking a data set to its publication is what an experienced analyst would do; what was missing was the disclosure of where the clue came from. Across the 30 audited sessions (Figure S2), no session read a file outside its own folder, but whether an instruction about memory was followed depended on its wording: a general instruction was broken in 12 of 16 sessions and an explicit one was kept in eight of eight. Neither reached the index, which was present in all 30 sessions, including the six sponge sessions (one of them interrupted) run with an agent type that had no memory tools. An instruction in the prompt is therefore not a boundary. Blind tests and analyses of unpublished data need technical limits: memory switched off or a separate account, a workspace without web access when the data must not be linked to outside sources, and an audit of every tool call (Figure S2c).

#### Practical limits

The workspace had about 8 GiB of RAM, which forced database slicing, disabled mass recalibration in FragPipe, and left the impurity correction out of the FragPipe TMT run. Slicing did not change identifications, because false discovery was estimated on the combined results; without recalibration FragPipe may have lost some identifications; and the FragPipe TMT output was not used for the main quantitative results; it served to compare identifications and, in the Dravet study, as a check on fold changes, which it compressed. Sessions were interrupted, and long jobs needed checkpointing and supervision (DogWatch). Copying multi-gigabyte RAW files from the researcher’s computer was slow, and the system package manager and conda were blocked, so tools had to be obtained as self-contained releases. Commercial software (PEAKS, Proteome Discoverer) could not run in the environment, so comparisons with those tools depend on deposited results or on a person running them with the same database and parameters. The agent wrote code, figures, and draft text but cannot take responsibility for them; the authors checked the results and are accountable for them.

### Relation to Other Agent Systems

Two recent systems are closest to this work (Table 8). In the first, one investigator built a complete proteomics analysis application from four natural-language prompts in under 10 min; the application normalizes, tests, and plots quantitative tables, and it reproduced outlier exclusion, PCA separation, and volcano plots of two published data sets.^3^ In the second, agents identify published data sets in the literature, download them, and requantify them with containerized search and quantification tools; the five worked examples needed human guidance, and the requantified results agreed with the deposited ones at per-sample correlations of 0.85–0.997 and differential-set overlaps of 0.44–0.51.^4^ Both start from a defined task, a tool to build or a published analysis to repeat, and judge success by agreement with a known result. Here the task was not fixed in advance: the questions arose from the data of unpublished studies, starting from RAW files, and success was judged by whether an answer changed what a study could claim. Such questions changed a database, a sample map, a statistical model, and a deamidation claim, and a question first raised by a simulated non-specialist, then asked by the researcher of every data set, exposed failed or partial alkylation in all three studies (Tables 3 and 6). The three approaches share a dependence on people: both reports and ours describe human guidance as necessary, and none measured how often the decisive questions would be asked without an expert; our simulated non-specialist test only bounds part of that dependence. What this work does not provide, and the second system does, is scale: one public reanalysis rather than a set of published studies, and no automated retrieval.

**Table 8.** This Work Compared with the Two Closest Agent Systems for Omics Data Analysis.

| Feature | Vibe-coded application<sup>3</sup> | Omics data discovery agents<sup>4</sup> | This work |
| --- | --- | --- | --- |
| **Aim** | Build an analysis application from natural-language prompts | Find, retrieve, and requantify published omics data; synthesize across studies | Answer questions about the researcher's own data sets as they arise |
| **Input** | Protein-level quantitative tables | Articles and repository data (DDA and DIA proteomics, RNA-seq) | RAW files of three unpublished studies and one public data set |
| **Who sets the task** | Investigator, before building | Predefined retrieval and reanalysis workflow | Researcher, one question at a time, after each result |
| **Human role** | Prompts, correction of one error, comparison with published results | Guidance in every reanalysis (software versions, normalization decisions) | Questions, suspicion of results, decisions about samples with the data owners |
| **Agent output** | One application (about 1,400 lines) | Requantified data, cross-study comparisons, meta-analysis | New code for each question, dated reports, eight checks, and a run supervisor |
| **Validation against known results** | Reproduction of published outlier exclusion, PCA, and volcano plots in two data sets | Per-sample $r$ 0.85–0.997 and differential-set overlap 0.44–0.51 against deposited results; data set identification precision 80% | Same peptide as Proteome Discoverer (99.2%) and FragPipe (99.3%); entrapment FDP 1.3%; three questions re-asked in six sessions |
| **Known defects and null tests** | Not part of the design | Informal security tests (injected prompt, fake identifiers) | 306 seeded channel swaps, 200 permutation nulls, seeded database variants |
| **Problems found in the analyzed data** | Not part of the design | Reporting gaps (for example, unlabeled sample columns) | Database lacking one symbiont, swapped channels, inflated statistics, false deamidations, failed alkylation |
| **Scale** | Two data sets | Five worked reanalyses and a literature corpus | Four data sets, one of them public |
| **Model and code** | Replit agent (model not named); application code on GitHub | LLM agents with tools served through MCP; code on GitHub, container images not distributed | Claude in claude.ai; Interroga package; three modules available on request |
| **Degree of autonomy** | Agent writes the application; the investigator tests it | Agents run retrieval and reanalysis; human guidance in each case | Agent acts only on the researcher's questions; decisions about samples are taken by people |
| **New analyses created during the work** | No; the application performs a fixed set of analyses | Within the predefined tools served to the agents | Yes; a new check for each question (Table 2) |
| Record of decisions | Prompts and application code | Code and evaluation scripts | Dated reports with code, inputs, answer, and the reason for each change |

## CONCLUSIONS

Conversational interrogation lets a researcher formulate and test analytical questions that the initial pipeline did not anticipate; in these four data sets, such questions revealed identification, quantification, and sample-quality problems that changed what the studies could claim. An LLM agent working in a conversation can also carry a proteomics analysis from RAW files to figures within a consumer subscription: a public TMT data set was reanalyzed in 24 min from one instruction and agreed with the established workflow at the level of spectra, fold changes, and the sample farthest from its group, and three questions, two on that data set and one on the mosquito study, gave the same answers when re-asked in new sessions of one model version; we did not compare this time with an analyst working without an agent. Its value, however, lies less in running standard steps than in making the next question cheap to ask. In the case studies, that changed a database, a sample map, a statistical model, and a deamidation claim. None of these problems was flagged by the pipeline as first run: a search database missing one of two symbionts, a swapped TMT channel, deamidation produced by the instrument’s choice of precursor peak, and, once the same question was asked of every data set, failed or partial cysteine alkylation in all three studies. Each was present in data already acquired and remained unexamined until a specific question prompted a closer look. The approach works as a loop between the people who generate the samples, the data, the analysis, and the biology, with a control in each test; the agent shortens each turn of that loop but replaces none of its parts. In tests with defects we seeded, the checks it produced detected the seeded defects of the class they were written for, of which standard sample QC detected 6–100%, and missed the others, including a 10% loss of one proteome that fell below DBScope’s 90% threshold by design; this does not establish superiority over sample QC beyond those defects. The evidence comes from four data sets, three of them our own, analyzed with the Claude models available in October 2026, and the agent tests used a single model version; testing other models, laboratories, and users on tasks defined in advance is the next step.

The main risks are the agent’s own analytical errors, metadata that are wrong, the need for a researcher who knows which questions to ask, and context stored in the researcher’s own account, which an agent session can draw on without being asked to. We recommend six practices: (1) keep each analysis version with its date and the reason for each change; (2) require that every reported number be regenerated from code, not recalled; (3) build controls into each new analysis; (4) supervise long runs with a standing watchdog rather than on request; (5) leave decisions about samples to the people who generated them; and (6) report whether stored memory was on, enforcing blind tests with technical limits (memory off or a separate account, restricted web access) rather than with instructions. The checks are released as tested code so that they can run without an agent.

## ASSOCIATED CONTENT

### Supporting Information

The Supporting Information is available free of charge at [link added by the publisher].

Supporting Tables S1–S14: question log by date of the dated report (S1); search, quantification, and statistics parameters (S2); error log (S3); inventory of scripts (S4); seeded channel swaps (S5); permutation null runs (S6); seeded database variants (S7); reanalysis of PXD063185 and comparison with the deposited results, with timings, reproducibility checksums, entrapment and phosphopeptide searches, and the independent agent sessions (S8); fragment-tolerance test of the Dravet Exploris 480 run (S9); researcher time by stage from file timestamps (S10); simulated non-specialist sessions, audit, and outcomes (S11); cysteine forms in the 54 sponge runs (S12); independence and status of the evidence for each claim (S13); map from each central result to the script and inputs that produce it (S14) (XLSX)

Supporting Figures S1–S2: fate of the MS2 spectra and open search in the mosquito holobiont (S1); use of the corresponding author’s stored memory in the 30 audited agent sessions (S2) (PDF)

Interroga package: tested modules, the laboratory pipeline, scripts as run, unit tests, and scripts that regenerate every figure and analysis from source data (ZIP)

Conversation transcripts with timestamps (PDF)

## AUTHOR INFORMATION

### Author Contributions

C.S.A.: investigation (sponge sample preparation and LC–MS/MS), data curation, writing – review and editing. M.K.: investigation (mosquito rearing, TMT sample preparation, and LC– MS/MS), data curation, writing – review and editing. A.C.S.R.: investigation (neurosphere culture, hyperthermia, and TMT sample preparation), data curation, writing – review and editing. A.L.V.: resources and discussion of the sponge study (natural products context), writing – review and editing. R.M.-d.-F.: conceptualization and supervision of the mosquito study, resources (mosquito lines), funding acquisition, writing – review and editing. M.Z.P.G.: conceptualization and supervision of the neurosphere study, resources (iPSC lines and neurospheres), funding acquisition, writing – review and editing. M.J.: conceptualization, methodology, formal analysis and software (through the AI agent), supervision, funding acquisition, writing – original draft.

All authors approved the final version.

### Notes

The authors declare no competing financial interest.

## ACKNOWLEDGMENTS

The authors thank the members of the mosquito, Dravet neurosphere, and sponge teams who contributed to the companion studies whose data served as case studies here, Joana Sandes for the sponge specimens and their taxonomic identification, and, in memoriam, Guilherme Muricy, who collected the V. ophiraphidites specimens. The biological results of these studies are reserved for the companion studies. We thank the authors of PXD063185 for the public data. This work was supported by CNPq (Research Productivity fellowship) and FAPERJ (Edital N° 22/2025 – Pensa Rio, grant E-26/210.840/2025, coordinated by A.L.V.; Temáticos E-26/211.348/2021). Use of AI: Claude (Anthropic), used through claude.ai in Project mode between 9 and 20 October 2026, wrote and ran all analysis code, built the eight tools in Table 2, drew the figures, and produced the first draft of this text in response to the corresponding author’s questions; the corresponding author reviewed every result and made decisions about samples together with the data owners, and all authors edited and approved the text.

## TABLE OF CONTENTS GRAPHIC (For Table of Contents Only)

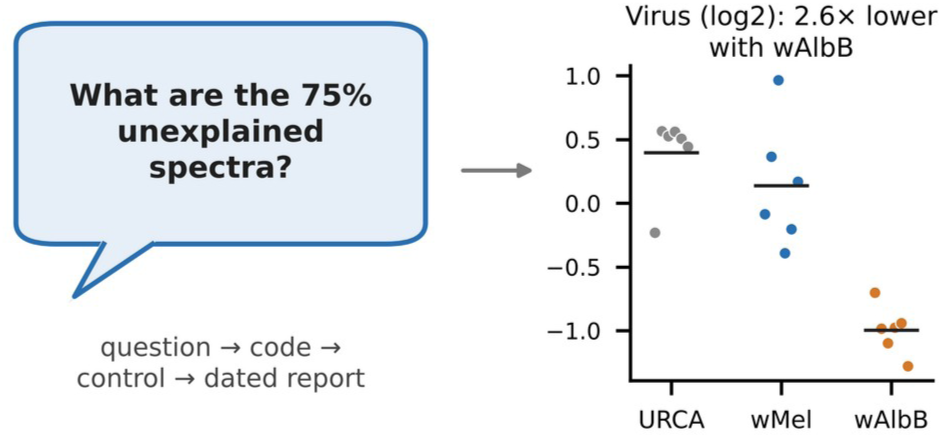

